# Repetitive mild traumatic brain injury causes neuronal damage in the APP/PS1 mouse model of Alzheimer’s disease without an enduring impact on amyloid pathology, sleep, or epileptiform activity

**DOI:** 10.1101/2025.06.06.658157

**Authors:** Jefferey Yue, Victoria Carriquiriborde, Wai Hang Cheng, Taha Yildirim, Jianjia Fan, Sean Tok, Michael Kelly, Cheryl L. Wellington, Brianne A. Kent

## Abstract

Traumatic Brain Injury (TBI) is a known risk factor for Alzheimer’s disease and related neurodegenerative diseases. Sleep disturbances and epileptiform abnormalities can appear after TBI and may contribute to the development of neuropathology. In this study, we characterized sleep, epileptiform activity, and neuropathology after repetitive mild traumatic brain injury (rmTBI) in a mouse model of Alzheimer’s disease. We used the Closed Head Impact Model of Engineered Rotational Acceleration (CHIMERA) to deliver rmTBI or sham (control) treatment to 6-month-old APP/PS1 mice (N=19). One month post-injury, we implanted electroencephalogram (EEG) and electromyographic (EMG) electrodes, recorded for 72 hours, and then collected brain tissue and blood plasma. Our assessment of sleep architecture showed that time spent in vigilance state was not affected by the rmTBI one month post-injury; however, power spectra analysis showed a shift towards higher frequencies in the rmTBI group during non-rapid eye movement (NREM) sleep. Epileptiform activity did not differ between sham and rmTBI. Compared to sham controls, the rmTBI group showed higher neurofilament light (NF-L), but not glial-fibrillary acidic protein (GFAP) in blood plasma and no change in Aβ pathology. These results indicate sustained neurological injury in the APP/PS1 mice one month after rmTBI without affecting amyloid deposition in the brain. Our study suggests that rmTBI can induce neural injury without causing enduring sleep disruption, seizures, and exacerbation of amyloidosis in the APP/PS1 mouse model.

## Introduction

Traumatic brain injury (TBI) is a leading cause of global morbidity and mortality and a risk factor for neurodegenerative diseases, such as Alzheimer’s disease (AD) ^1,2^. TBI severity ranges from mild to severe, with mild TBI (mTBI) accounting for approximately 90% of cases ^3^. In people age 65 and older, TBI account for 80,000 emergency hospital visits each year in the US, with falls being the primary cause ^4^. It has also been shown that individuals who experience a TBI in this age group (65+) have an increased risk of developing dementia ^5,6^. Additionally, it is thought that repetitive mTBI (rmTBI) synergize and have an even greater impact on cognitive and executive function ^7,8^. This is why it is imperative to understand the changes that rmTBI causes in brain activity and how this is associated with late-life neurodegeneration and AD development.

Pre-clinical studies using animal models have shown that rmTBI are associated with increased neuronal excitability, neurological, motor, and cognitive impairments, and axonal damage within the first two weeks after injury ^9–11^, all of which are hallmarks of AD pathology. Some of the behavioral and neuropathological outcomes of rmTBI have also been reported at chronic timepoints, up to 12 months post-TBI ^10,12^. rmTBI in transgenic animal models of AD have also shown exacerbated neuropathology development, including the accumulation of Aβ, at both acute and chronic timepoints post-injury ^13–15^.

In addition to impacting cognition and increasing the risk of neurodegenerative disease, rmTBI can cause sleep disturbances and posttraumatic epilepsy (PTE) ^9,16–18^, both of which are independent risk factors for neurodegenerative disease ^19–21^. The most prevalent sleep disturbances reported in humans after rmTBI are insomnia and hypersomnia ^17,22^. In mouse models, rmTBI can cause reduced sleep duration, up to 16 months post-injury ^23^, as well as increased sleep fragmentation ^16^. Other EEG measures, such as spectral power, have also been shown to be affected by rmTBI ^24–26^. Spectral power is a quantitative EEG analysis that measures the strength of the brain signal at each frequency. Different spectral power patterns have been associated with distinct mental states and brain functions ^27–30^, which makes spectral power a useful tool to compare healthy and diseased brain states. One of these functions is glymphatic clearance, the system that promotes the clearance of metabolic waste products such as Aβ and tau ^31–33^. Glymphatic function has been positively correlated with low-frequency power (delta band) and negatively correlated with high-frequency power (beta band), both in wild-type mice ^34^ and in humans ^35^. Therefore, shifts in spectral power could be a potential mechanism linking rmTBIs and clearance of pathological proteins driving AD.

Sleep disturbances after rmTBI can also be accompanied by PTE in humans ^36,37^. The proportion of patients with PTE increases with injury severity ^37^, yet it is still unclear whether repetitive impacts can increase PTE susceptibility. How the post-rmTBI sleep disturbances or epileptiform activity contribute to the risk or progression of neurodegeneration is not yet clear.

Here, we investigated the effects of rmTBI in APP/PS1 mice ^38^, by EEG/EMG recording and quantification of multiple protein targets through immunoassays. As most previous studies have focused either on severe TBI or acute time points post-injury, to our knowledge, this is the first study investigating the mutual effects on sleep impairment, epileptiform activity, and neuropathology after rmTBI in a mouse model of AD one month post-injury. We hypothesized that one month post rmTBI, mice would exhibit sleep impairments, shifts in spectral power, and increased epileptiform activity, plaque burden, and neuronal damage. We found that even though there were markers of enduring neuronal damage, there were no changes in sleep architecture, seizure susceptibility, epileptiform activity, or amyloid pathology one month post-injury. However, rmTBI was associated with an exacerbated spectral power shift relative to sham treated mice, suggesting a neuropathological mechanism that may accelerate the progression of AD.

## Methods

### Animals

All procedures in this study were approved by the Simon Fraser University Animal Ethics Committee (Protocol 1330P-21) and followed the guidelines from the Canadian Council on Animal Care. Figure 1 illustrates the study protocol. This study adheres to the Animal Research: Reporting of In Vivo Experiments (ARRIVE) guidelines for reporting animal research.

**Figure 1.**
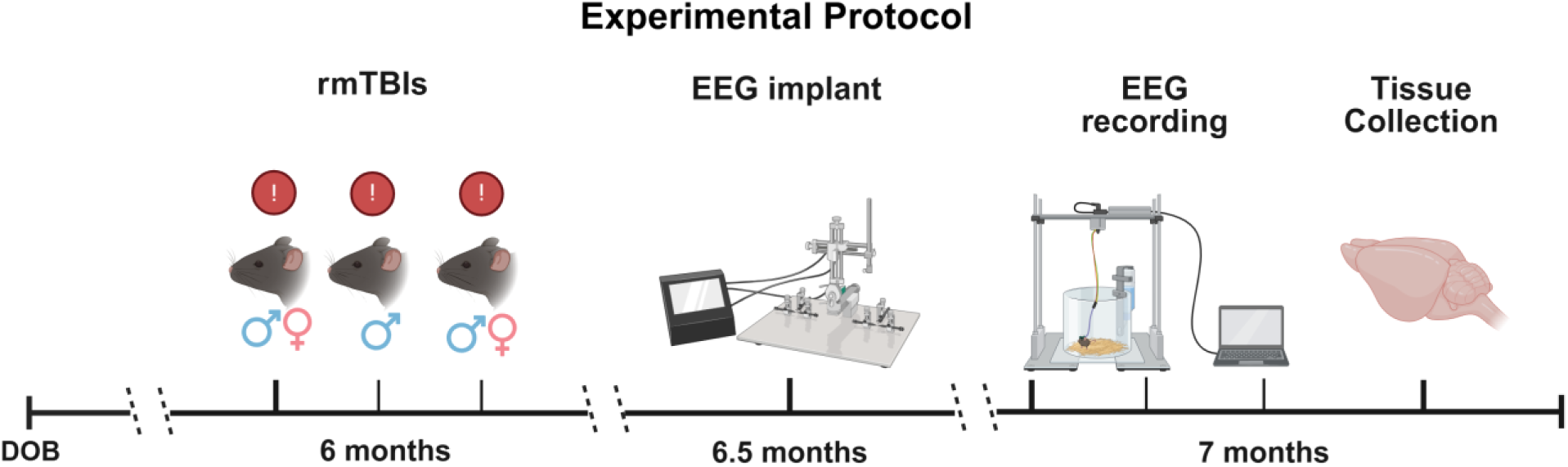
Schematic diagram illustrating the experimental protocol. At six months of age APP/PS1 mice received rmTBI. Male mice received 3 consecutive mTBI 24 h apart, and female mice received two mTBI 48 h apart. After recovering, mice received EEG/EMG intracranial implantation. Two weeks after implantation, at 7 months of age, the mice underwent 72 h of EEG/EMG recording. Tissue collection was completed immediately after recording. Created in BioRender https://BioRender.com/6h4as63

The mouse strain used for this research project, B6C3-Tg(APPswe,PSEN1dE9)85Dbo/Mmjax, RRID:MMRRC_034829-JAX, was obtained from the Mutant Mouse Resource and Research Center (MMRRC) at The Jackson Laboratory, an NIH-funded strain repository, and was donated to the MMRRC by David Borchelt, Ph.D., McKnight Brain Institute, University of Florida. APP/PS1 mice are double transgenic mice that overexpress mouse/human APP containing the Swedish mutation (K595N/M596/L) and a mutant human presenilin 1 (PS1 delta E9). This mouse line was selected because it had been previously studied using CHIMERA TBI ^12^ and EEG assessment of sleep and power spectra ^39^. Others have also reported epileptiform activity in this mouse line ^40,41^. This study aims to examine these outcomes concurrently in the context of TBI for the first time. Both males (n=8) and females (n=11) were used in this study. All 19 animals were single housed in plastic clear cages under a 12:12 LD cycle. Food and water were available *ad libitum*.

### 1. Repetitive mild TBI (CHIMERA)

CHIMERA procedures followed the protocol previously described ^11,12,42^. CHIMERA produces a diffused brain injury (Namjoshi et al., 2014). The piston impact is combined with free head movement, resulting in a rotational acceleration-deceleration mimicking the mechanisms underlying diffused axonal injury. This feature was the reason for selecting the CHIMERA system for the present study, as it more closely replicates the biomechanics of the most common forms of human TBI. APP/PS1 mice at 6 months of age were randomly assigned to sham (n=8) or rmTBI (n=11) groups. TBI was administered at 6 months of age, corresponding to the onset of amyloid plaque deposition and rapid amyloid plaque growth in APP/PS1 mice^43,44^. This time point is widely used in preclinical AD research because it models early pathological progression, analogous to the prodromal phase of AD in humans, enhancing clinical relevance. Mice received meloxicam subcutaneously (SQ, 1mg/kg) 30min before being restrained on the CHIMERA device. Mice were anesthetized with isoflurane, received saline (SQ, 0.5mL) for hydration, and eye gel was applied. Total duration of isoflurane exposure was 4-8 min. Sham animals underwent the same procedures except for the CHIMERA impact. For the rmTBI group, the intensity of the impact was lowered for females to compensate for lower body weight (weight mean ±SEM: male 35.2g ±1.26, female 27.1g ±1.03) and to reduce morbidity. Male mice received 3 mTBI (direct impact at 0.7J) 24 hours apart, and female mice received 2 mTBI (direct impact at 0.6J) 48 hours apart. The 0.7J impact energy is the highest energy within the mTBI parameter as previously defined ^42^. The positioning of the head was as described in Namjoshi et al. (2014). The piston struck the head in an area of 5mm surrounding bregma, making it a bilateral injury. Standardized procedures were used to ensure consistent head positioning across injuries and animals. Females were subjected to a modified rmTBI protocol based on findings from a pilot study. During the pilot study, two female mice exposed to the 3 mTBI at 0.7J protocol exhibited severe paralysis following the second and third injuries, leading to mortality after the final injury. We concluded that the differential response may be attributable to baseline weight differences between male and female mice (weight mean ±SEM: male 33.0g ±0.9, female 26.4g ±0.8). To ensure that female mice sustained a brain injury consistent with the expected outcomes of mTBI, females were only administered 2 rmTBI in this experiment. After procedures, mice recovered in a warmed cage supplemented with hydrogel and were returned to the housing room two days after the final mTBI.

### 2. Injury effects: LRR and Grimace

The loss of righting reflex (LRR) was calculated as the time interval from the termination of isoflurane to the successful attempt of righting for each mouse after injury. The Grimace Scale is a two-point scoring system based on defined facial and physical features of the mice as indicators of pain and discomfort ^45^. Grimace score was recorded once every five minutes. The Grimace recovery time was calculated as the time interval between when each mouse righted their body and the time when all grimace scores were at zero.

### 3. Electrode implantation

2EEG/1EMG headmount (catalog number 8201-SS; Pinnacle Technology, Lawrence, KS, USA) implant surgery was performed as previously reported ^39,46^. Mice were anesthetized with isoflurane and given meloxicam intraperitoneally (IP, 5 mg/kg), lidocaine (SQ, 7 mg/kg), and lubricating eye gel. Total duration of isoflurane exposure was 40-50 min. A middle rostral-caudal incision was made to expose the skull surface. Four stainless steel screw electrodes were inserted into the skull at coordinates relative to bregma: AP +/- 3mm, ML +/- 1.5mm. EMG wires were inserted into the nuchal muscle. A two-part epoxy was applied to the screws to ensure good electrical contact, and a multimeter was used to test the electrical contact. The headmount was secured with a layer of dental cement (catalog number 525000; A-M Systems, Sequim, WA, USA). Post-operatively, lactated ringer (SQ, 5mg/kg) was provided for hydration, and buprenorphine (0.05mg/kg) was provided for 3 days for pain relief. All mice were allowed to recover for at least 10 days before EEG recordings.

### 4. EEG recording

EEG/EMG recordings were collected in a tethered system (8200-K1, Pinnacle Technology) using Sirenia® Acquisition software (version 2.2.1). Sample rate was 400Hz, preamp gain was 100dB, and low-pass filter was 40Hz for EEG and 100Hz for EMG. EEG/EMG was recorded for 72 hours.

### 5. Assessment of vigilance state

Assessment of vigilance state was performed using the final 24 hours of the EEG recording, allowing for 48 hours of habituation. An experimenter blinded to the experimental condition used cluster scoring followed by visual inspection to score each 10s epoch for wake, NREM, and REM states using Sirenia Sleep Pro software (version 3.0.1). Wake state was defined by a low-amplitude EEG, with a dominant frequency higher than 4Hz, typically accompanied by high EMG activity. NREM was defined by high-amplitude EEG and a dominant frequency of less than 4Hz, usually accompanied by low EMG activity. REM was defined by low-amplitude and uniform waveforms with peak frequency at 4-8Hz and occurring following NREM state. The state of each epoch was defined based on the state dominating more than 50% of the epoch. Epochs containing artifacts (i.e., extremely high power) or without clear recordings (i.e., flat line) were excluded from the analyses. A bout was counted when the animal remained in the same vigilance state for 30s (3 epochs) or more. Bout length was calculated as the average time the animal spent across bouts of NREM, REM, or wake respectively.

### 6. Spectral Power Analysis

Spectral power provides a quantitative measure of how neural activity is distributed across the frequency bands of interest. We calculated power at each frequency and each electrode separately. First, the raw data were pre-processed by splitting the signal into 10s epochs and applying a 2^nd^ order Butterworth filter (0-40Hz). Then, Welch’s method was used to transform the filtered data into the frequency domain using a Hamming window of 10s with a 50% overlap. To calculate relative power, absolute power at each frequency was divided by the total power across frequencies (0.5-30Hz), comprising the delta (0.5-4Hz), theta (4-8Hz), alpha (8-13Hz), and beta (13-30Hz) bands previously defined in the literature ^34,39,47^. We only included frequencies 0.5-30Hz to avoid distortions caused by the filter cut-off. We used the PLS method to analyze spectral power data. The input data was a matrix wherein rows represented each individual mouse separated by treatment group (TBI vs sham), and the columns represented the average power at each frequency (0.5-30Hz).

### 7. Epileptiform Activity Analysis

Seizure detection was performed manually using the Sirenia Seizure Pro software (version 3.0.1). Two researchers blinded to the experimental condition visually inspected the 72 hours recordings for seizures. Seizures were defined as high-amplitude and high-frequency events more than 10s in duration. Heat maps and spectral plots were used to identify epochs with high EEG power and high frequency spectra to aid in seizure detection. Once a seizure was identified, it was marked, and the window size, step size, and power measurements from this reference seizure were used to identify other seizures.

To detect epileptiform activity (EA) events, we used a semi-automated method to score the last 24 hours of the recording. We used a bandpass filter (0.5-40Hz) to filter out the high-frequency artefacts and visually identified an example of a spike, poly-spike, and spike-wave discharges (SWD) that fulfilled the definition of each category according to the recommendations by Jin et al.^48^. A spike was defined as a cortical epileptiform spike (wave) that showed more than 2x the baseline voltage mean. Poly-spikes were defined as two or more spikes with the same amplitude criteria as a spike occurring in succession. Lastly, a SWD was defined as surface-negative spikes alternating with surface-positive waves or vice versa at 7-10Hz with durations between 0.5-5s. All EA were visually verified by a researcher blinded to treatment.

### 8. Biosample collection

Mice were anesthetized using isoflurane and then euthanized by CO2. Blood samples were collected via cardiac puncture and mixed with 100mg/mL EDTA to prevent coagulation. Blood samples were centrifuged at 1500g for 20min, and supernatant was collected as plasma. Plasma samples were stored at -80°C until analysis. Following cardiac puncture, mice were transcardially perfused with ice-cold phosphate buffered saline (PBS, pH 7.4), and brain tissue was extracted. Since injuries were delivered at the midline along the sagittal axis of the brain, we expect similar injury in both hemispheres; hemibrains were randomly sampled for differential processing prior to biochemical and histological analyses. One hemibrain was frozen on dry ice then stored at -80°C, and the other hemibrain was fixed in 4% paraformaldehyde (PFA) for 3 days, which was then cryoprotected in 30% sucrose. The cryopreserved hemibrain was coronally sectioned at 40μm using cryotome (Leica).

Hemibrain lysate was generated in serial extraction as described with modification ^13^. Frozen hemibrain was ground using a mortar and pestle, while keeping it at a low temperature with liquid nitrogen. Ground tissue was then homogenized (Ultra-Turrax T25) in four volumes of carbonic buffer containing 100mM Na_2_CO_3_, 50mM NaCl, and 1mM phenylmethanesulfonyl fluoride (pH 11.5), supplemented with mini-cOmplete protease inhibitor cocktail (1 tablet/10 ml; Roche, cat. # 11836170001) for 20s, then sonicated in 20% output for 10s. The samples were then centrifuged at 16000 g for 45min. The supernatant was collected and neutralized in 1.5 times the volume of 1M Tris-HCl buffer (pH 6.8) to yield a final pH of ∼7.4. The pellet was resuspended in four volumes of 5M guanidine hydrochloride (5M GuHCl, 50 mM Tris-HCl, pH 8.0) and incubated at room temperature with rotation for 3 hours. Both fractions were stored at -80°C until analysis.

### 9. Plasma Biomarker Analysis

Plasma levels of murine GFAP were quantified by a customized immunoassay using a pair of rabbit antibodies against mouse GFAP (Abcam, cat # ab244094) with the MesoScale Discovery (MSD) platform as previously published ^49^. Plasma was diluted 4 times with PBS containing 1% bovine serum albumin (Sigma Aldrich, cat. # A7906). Plasma levels of murine Nf-L were measured by the MSD Neurofilament-light assay kit (Mesoscale, cat. # K1517XR-2) following the manufacturer’s instructions. Murine plasma was diluted 6 times with MSD Diluent 12 for the Nf-L assay. The researcher conducting biomarker analysis was blinded to the experimental conditions. All samples with reported comparisons were analyzed within the same run.

### 10. Immunohistochemistry

Immunohistochemistry of amyloid plaques was performed with the regions of interest (ROI) defined according to anatomical landmarks illustrated on the Allen mouse brain atlas (https://mouse.brain-map.org/static/atlas). We selected brain regions that are reported to exhibit early Aβ accumulation in AD mouse models, particularly following sleep fragmentation, as well as in individuals with early AD^50–52^. The ROIs include parietal isocortex (ISOCTX) and hippocampus (HPF). The parietal cortex was further divided into retrosplenial cortex (RSP), motor sensory cortex (MS), entorhinal cortex (ENT), and piriform area (PIR) for regional analysis. Anterior cortical regions investigated include the anterior cingulate area (ACA), frontal motor sensory cortex (F-MS), and frontal piriform area (F-PIR).

Every tenth coronal brain sections at 40 µm-thickness were sampled, and up to three sections per region were analyzed. The sections were washed in 0.05% Triton X-100 diluted in TBS (TBS-T), then incubated serially in 88% formic acid for 5 min, 0.3% hydrogen peroxide diluted in 0.25% Triton X-100, and 5% milk. Sections were then incubated with primary antibody (6E10: 1:1000, BioLegend cat. # 803004) overnight at 4°C with rocking motion. After washing in TBS-T, sections were incubated in biotinylated secondary antibody (1:1000, Southern Biotech, cat. # 1071-08) for 90 min in room temperature. After washing, the sections were developed with ABC reagent (1:400, Vector Laboratory, cat. # PK-6100), then stained in 0.2 mg/ml 3,3’-Diaminobenzidine (DAB). The sections were mounted onto SuperFrost microscope slides (FisherScientific. cat # 12-550-15) and coverslipped with DPX mounting medium (Sigma Aldrich cat. # 06522). The sections were imaged using an Axio Scan.Z1 slide scanner (Zeiss) at 20x magnification, followed by resizing at 30% and image optimization using the Zen software (version 3.7).

### 11. Aβ_42_ ELISA

Soluble (carbonic fraction) and insoluble (GuHCl fraction) Aβ_42_ in hemibrain lysates were quantified using a commercial ELISA kit (WAKO, cat # 296-64401) according to the manufacturer’s protocol. The carbonic fraction was diluted in 1:100-500, and the GuHCl fraction was diluted in 1:1000-5000. Every sample was loaded in triplicate, and the average value of the three readings was reported.

### 12. Statistical Analysis

The Shapiro-Wilk test was used to assess data normality ^53^. For LRR and Grimace scores, data were analyzed using repeated measures two-way ANOVA, followed by post hoc t-tests using the Šídák’s correction when appropriate. Male and female mice were analyzed separately.

Graphs and statistical analyses for the vigilance state data were performed using GraphPad (version 10.0.3) and RStudio (version 2024.04.1+748). Mann-Whitney was used to compare rmTBI and sham groups for duration, transition counts, bout counts, bout lengths, and lights on and off states for each vigilance state separately (NREM, REM, wake). Despite the adjusted injury protocol, males and females exhibited comparable levels of neural injury and injury-related behavioral effects. Therefore, data from both sexes were pooled for our main statistical analysis to account for the reduced sample size. However, we conducted a secondary analysis to evaluate sex differences using the Kruskal-Wallis test with Dunn’s correction for multiple comparisons. Normalized EEG spectral power data were analyzed using mean-centered partial least squares (PLS) analysis ^54^ in Matlab (MathWorks R2023a, version 9.14). PLS is a multivariate method adapted for neuroimaging data where dependent measures (e.g., power spectra frequencies) are highly correlated. We used PLS because it allowed us to detect the effects of rmTBI across the whole EEG frequency spectrum in a single analytical step, avoiding the need for multiple comparisons ^55^. The data structure and analysis steps were similar to those used in Cragg et al. ^55^ for power spectra analysis. Through this analysis, we obtained latent variables (LV) for each EEG channel (frontal and parietal) in each vigilance state, corresponding to the contrast and the pattern across frequencies. Statistical significance was established using permutation testing and bootstrap resampling for each LV.

All graphs and statistical analyses of epileptiform activity data were performed using GraphPad (version 10.0.3). Statistical significance for the non-parametric data was tested by Mann-Whitney to compare single spike, poly spikes, and SWD counts between rmTBI and sham groups. Total epileptiform activity across vigilance states was analyzed using Mann-Whitney multiple comparisons with Holm-Šídák correction. Similarly to the sleep assessment, sex differences in epileptiform activity were assessed in a secondary analysis using Kruskal-Wallis with Dunn’s correction for multiple comparisons.

Effects of rmTBI on plasma levels of Nf-L and GFAP were analyzed using Mann-Whitney with the assumption that sex does not affect baseline level of both plasma biomarkers. Amyloid histopathology was quantified using ImageJ by measuring the percentage of immunoreactive area within the defined ROIs. Statistical analyses of amyloid histopathology and Aβ_42_ ELISA assay were performed by analyzing main effects (injury vs sex) using two-way ANOVA on GraphPad. We used the two-way ANOVA for histology and ELISA analyses to assess sex differences because females have a greater amyloid burden compared to males when controlling for age at baseline in this mouse strain^56^.

Statistical significance was set at p≤0.05 for all analyses.

## Results

### rmTBI induced prolonged recovery in female mice

We first characterized injury responses by recording the latency of LRR and Grimace recovery after each mTBI. LRR is used in preclinical TBI studies as a proxy for loss of consciousness in humans, since LRR is associated with memory impairments and neurological abnormalities ^57^. Both sham and TBI mice experienced a similar duration of LRR (Fig. 2A & B). This is consistent with previous CHIMERA characterizations of very mild TBI injury where axonal injury and cognitive impairments were still present ^12^. As observed in the Cheng et al. ^12^ study, we also found a decrease in righting time on the second and third injury days. There was a main effect of injury on grimace recovery time in females (*p*=0.0099, F (1,9) =10.62) with longer delay in grimace recovery observed in the TBI group compared to the sham controls (Fig. 2D). However, there were no significant differences in the male group (*p*>0.05; Fig. 2C). No mice were withdrawn from the study due to morbidity or mortality. These data indicate that the TBI energy delivered to mice aligns within the mild intensity as previously reported ^42^.

**Figure 2.**
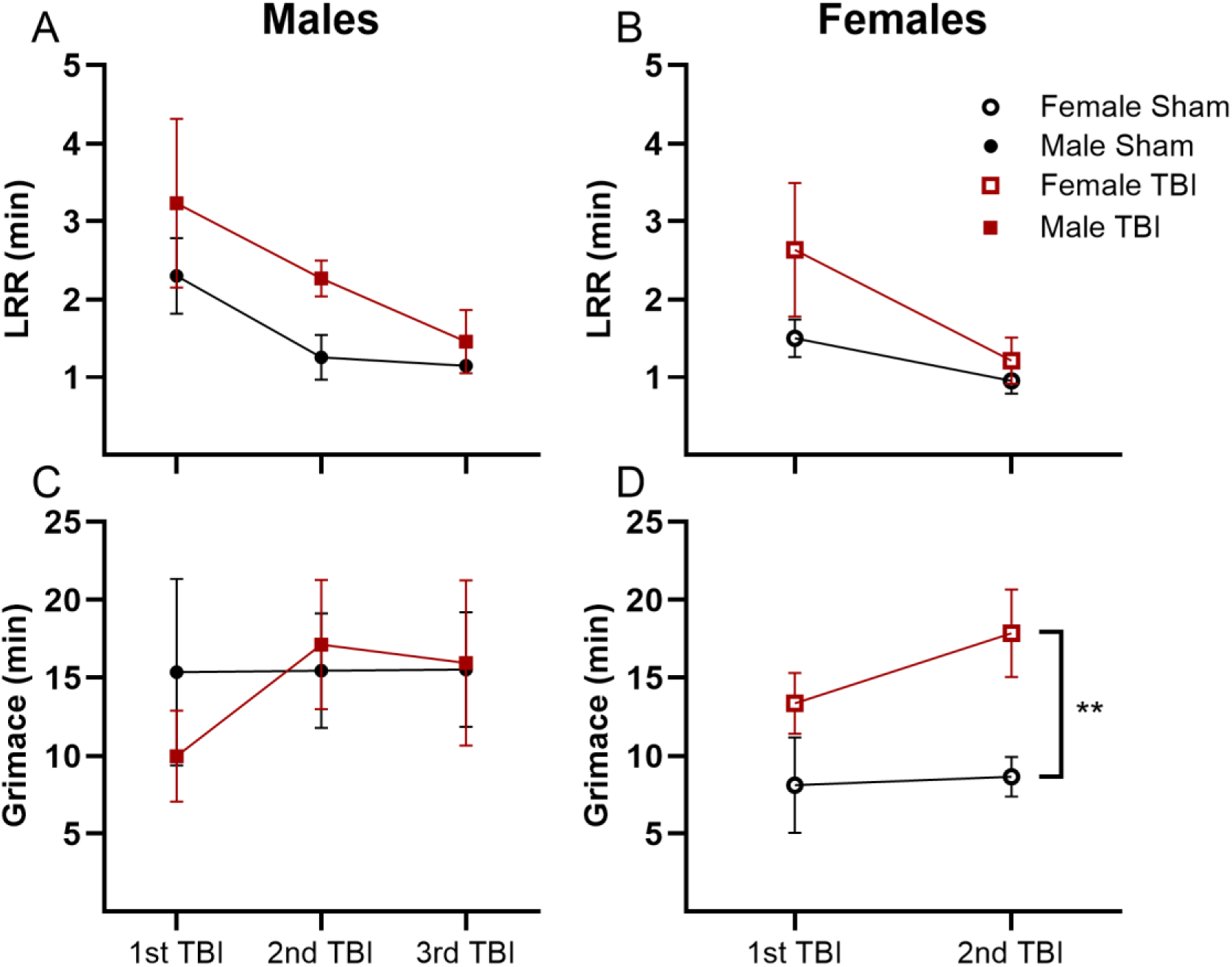
Neurological behaviour after rmTBI. Average LRR duration in minutes for each cohort after each injury timepoint for **(A)** males and **(B)** females. Average amount of time it took to reach a score of zero in the grimace scale in minutes after each injury for **(C)** males and **(D)** females. Results were reported as mean ± SEM and analyzed by repeated measures two-way ANOVA. Sham data is shown in black and rmTBI data is shown in red. Male TBI n=5, male sham n=3; female TBI n=6, females sham n=5. ** p< 0.01.

### Sleep architecture did not change one month post-injury

One female TBI mouse was excluded from all EEG analysis due to poor data quality (i.e., flat line, signal clipping, artifacts). Therefore, only 10 females (n=5 rmTBI, n=5 sham) were used for the EEG analysis. The percentage of time spent in NREM, REM, and wake was similar across the two groups. A Mann-Whitney test revealed no statistically significant difference between the rmTBI and sham groups for the vigilance states (*p*>0.05; Fig 3A). To evaluate the effects of rmTBI on sleep fragmentation, we assessed transition counts, bout counts, and bout lengths. There were no statistical differences between the rmTBI and sham groups in transitions between NREM-REM, NREM-wake, REM-NREM, REM-wake, wake-NREM states (*p*>0.05; Fig 3B). Bout counts and bout lengths for NREM, REM, and wake were not significantly different between the rmTBI and sham groups (p>0.05; Fig 3C & D). Sex effects for time spent on each vigilance state, transitions, bout counts, and bout length were analyzed separately with a post-hoc test. No statistically significant difference was found between males and females for either of the measures (*p*>0.05; Supplementary Fig S1). Additionally, we analyzed vigilance states separately during the 12 hours light phase and 12 hours dark phase. There were no significant differences between the TBI and sham groups in either of the states during the light or dark phases (*p*>0.05; Supplementary Fig. S2).

**Figure 3.**
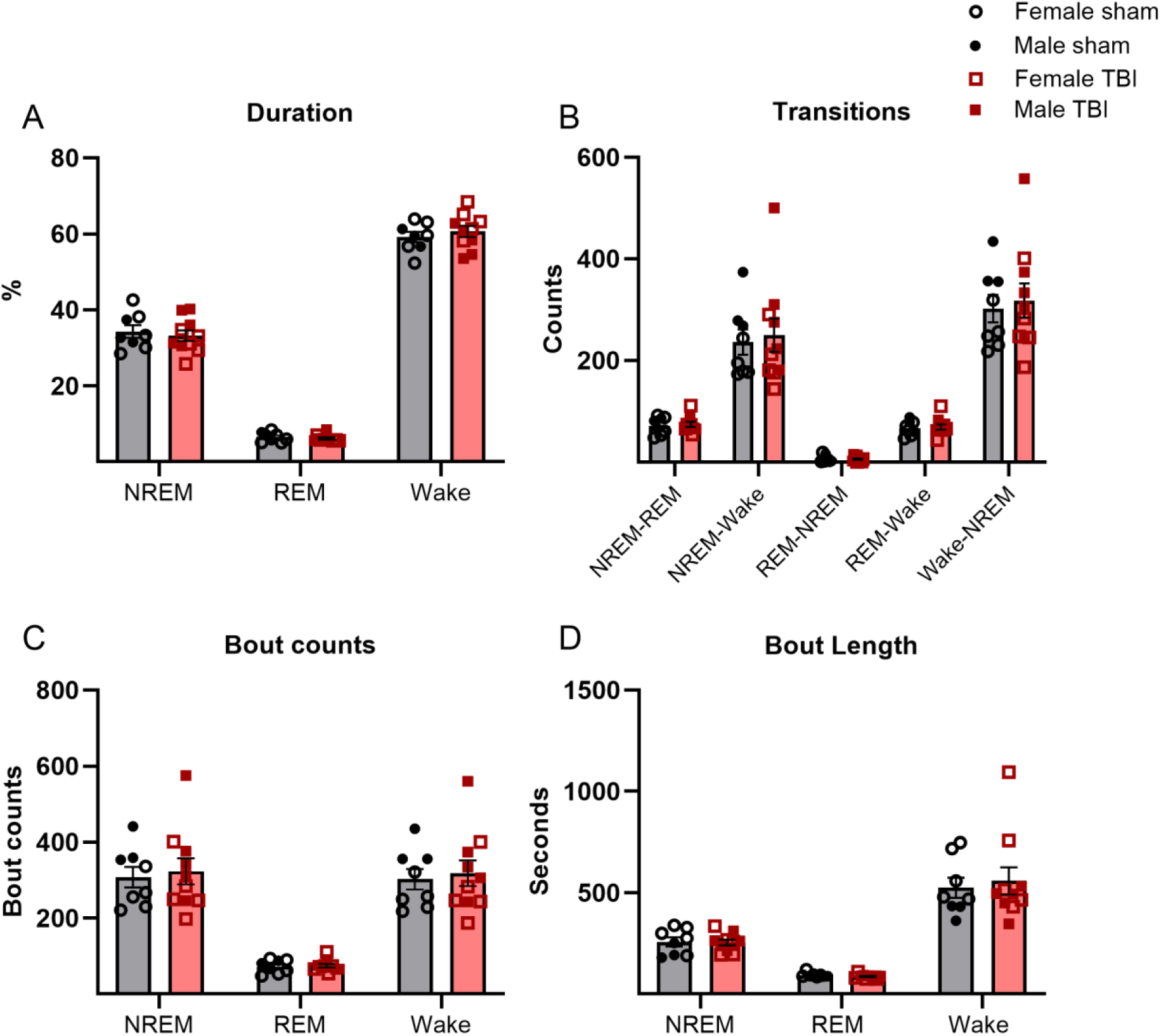
Sleep architecture one month after rmTBI in APP/PS1 mice. **(A)** Percentage of time spent in NREM, REM, and wake. **(B)** Transition counts between vigilance states NREM-REM, NREM-Wake, REM-NREM, REM-Wake, Wake-NREM. **(C)** Number of 30 s bouts of NREM, REM, and wake. **(D)** Average bout length. All data are expressed in mean ±SEM and analyzed by Mann Whitney-U test. Sham data are shown in black and rmTBI data are down in red. Females are shown with open symbols and males are shown with solid symbols. Each dot represents the average for a single mouse. Male TBI n=5, male sham n=3; female TBI n=5, female sham n=5.

### RmTBI caused spectral power shifts during NREM sleep

The PLS analysis identified a significant LV (*p*=0.041), indicating a difference between the TBI and sham groups during NREM. The TBI group exhibited higher relative power compared to the sham group at the parietal electrode at 5 Hz (theta band) during NREM (Fig. 4). Additionally, the TBI group exhibited higher relative power compared to the sham group across the higher frequencies (11-18 Hz) at the same electrode during NREM (Fig. 4). There was a similar trend at the frontal electrode during NREM, but it was not statistically significant (*p>*0.05; Supplementary Fig. S3 D). There were no differences observed between the TBI and sham groups during REM at either of the electrode locations (*p*>0.05; Supplementary Fig. S3 B & E). The spectral graphs during wake show some differences between the TBI and sham group (Supplementary Fig. S3 C & F) in the lower (4-6 Hz) and higher frequencies (20-24 Hz), but none of these reached statistical significance (Parietal EEG: *p*=0.069; Frontal EEG: *p=*0.0559) perhaps due to the low sample size.

**Figure 4.**
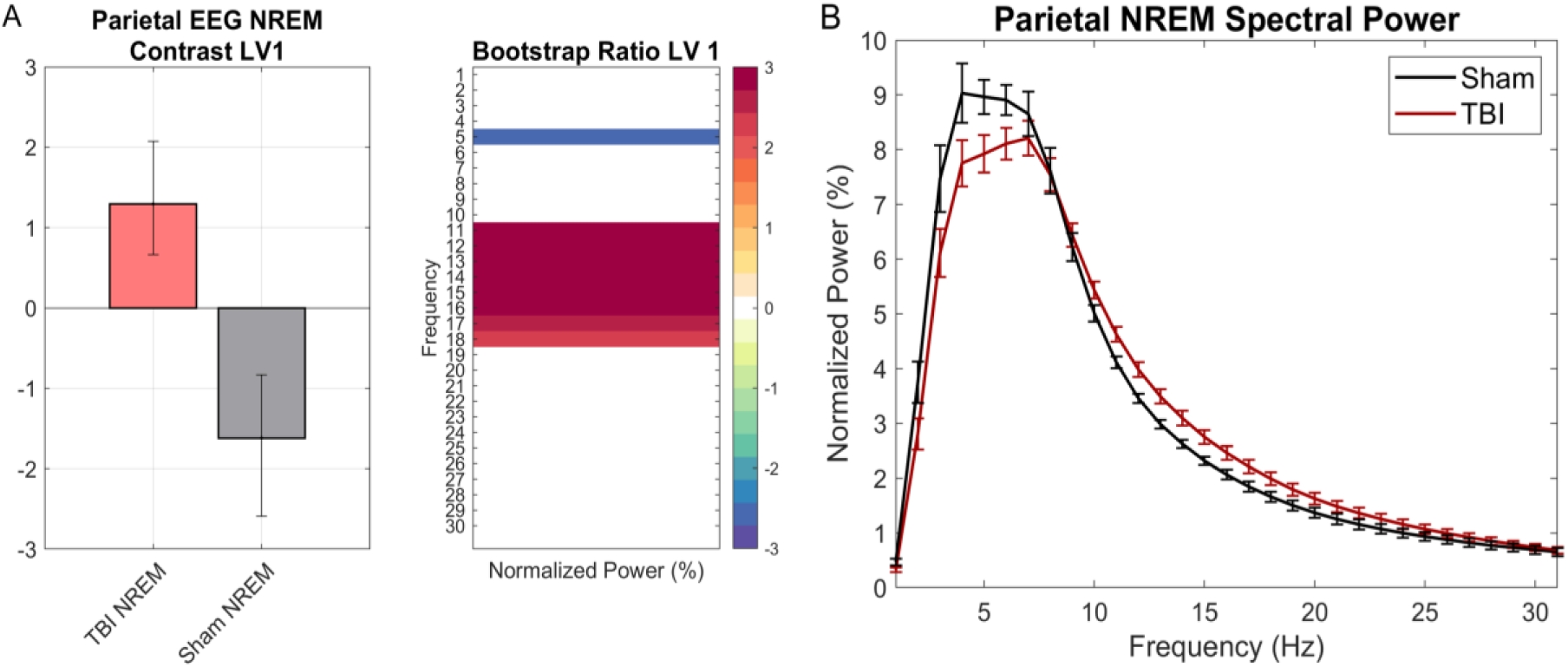
PLS analysis of the spectral power during NREM at the parietal EEG after rmTBI. **(A)** The PLS contrast in TBI and sham groups during NREM. Error bars represent the 95% confidence interval. The heat map represents the bootstrap ratio of the normalized power for each frequency. Red colors denote higher power in the TBI group whereas blue colors denote higher power in the sham group. Frequencies highlighted by these colors show statistical significance. **(B)** Spectral power with frequencies 0.5-30Hz in the x-axis and the normalized power values in the y-axis. Error bars represent the ±SEM. Sham data is shown in black and the rmTBI data is shown in red. Male TBI n=5, male sham n=3; female TBI n=5, female sham n=5.

### Epileptiform activity did not increase one month post-injury

Only one mouse exhibited spontaneous seizures (male, sham). Mann-Whitney revealed no statistical significance in Single Spike (SS), poly spikes (PS), or spike-wake discharges (SWD) counts per hour between the rmTBI and sham groups (*p*>0.05; Fig. 5A-C).

**Figure 5.**
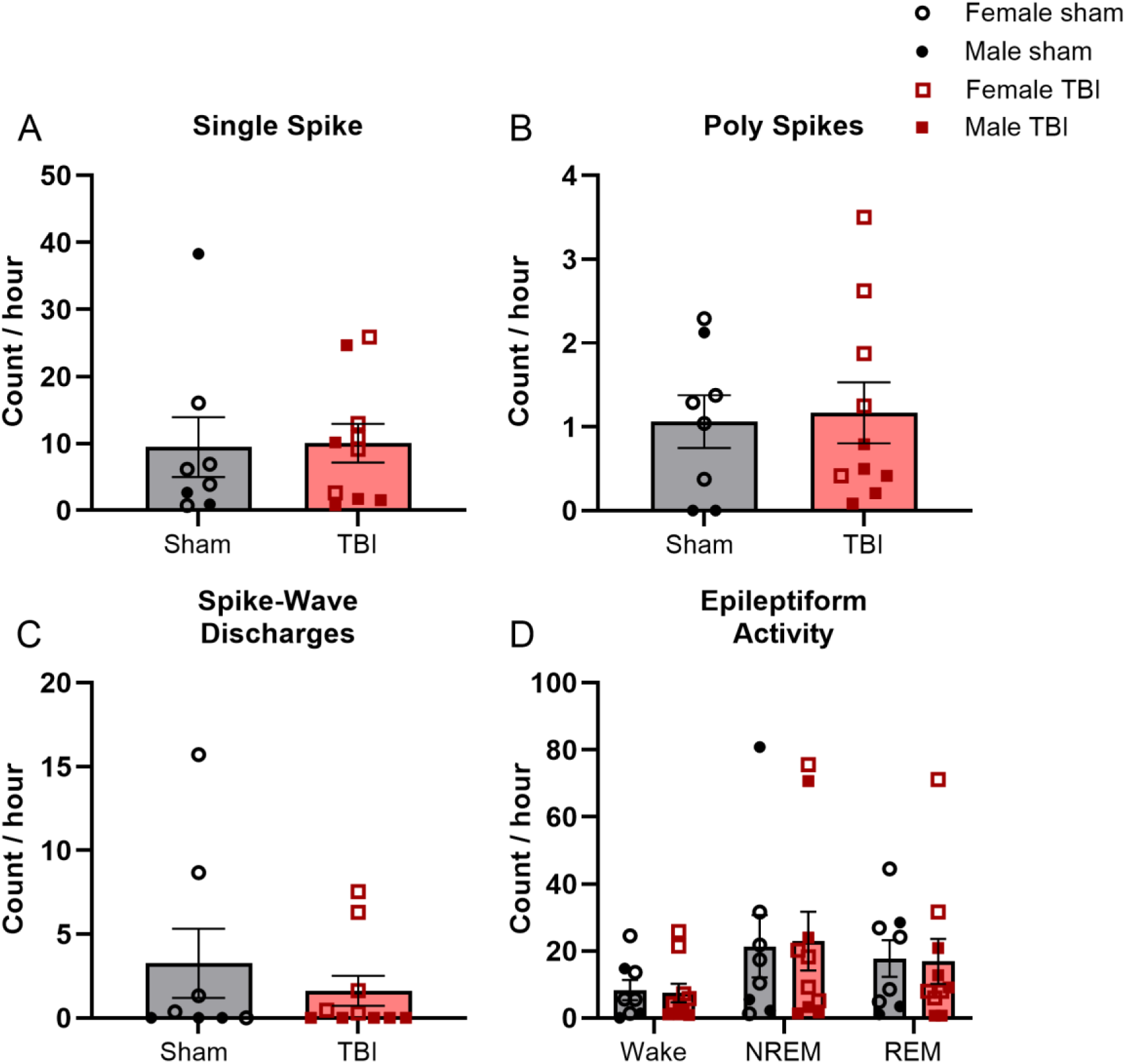
Epileptiform activity after rmTBI. **(A)** Single spike (SS) count per hour. **(B)** Poly-spikes (PS) count per hour. **(C)** Spike-Wave discharges (SWDs) count per hour. **(D)** Epileptiform activity (EA) count was obtained by adding SS, PS, and SWDs. Data are reported as mean ±SEM and analyzed by Mann Whitney-U test. Sham data are shown in black and rmTBI data are shown in red. Females are shown with open symbols and males are shown with solid symbols. Each data point represents the average for a single mouse. Male TBI n=5, male sham n=3; female TBI n=5, female sham n=5.

Additionally, we compared the prevalence of epileptiform activity during the three vigilance states and found no significant differences between the groups (Fig. 5D). However, as observed in previous studies ^40,58^, there was a trend towards an increase in epileptiform activity (EA) prevalence during the sleep states (NREM and REM).

Due to the small sample size, we did not include a statistical analysis of sex differences in our main analysis. However, SWDs were only observed in female mice in both groups. A post-hoc test showed that the difference in SWDs counts between males and females was statistically significant within the rmTBI group (*p*=0.016; Supplementary Fig. S4) but not the sham group (*p*=0.065). The post-hoc test also showed that females had significantly more EA per hour than males in the rmTBI group, but only during wake (*p*=0.025). The other two states, NREM and REM, had no significant sex differences for total EA per hour (*p*>0.05). There were no sex differences in singles spike or poly spikes counts (*p*>0.05).

### rmTBI induced neurological injury

We assessed the release of two cytoskeletal proteins, Nf-L from neurons and GFAP from astroglia, by performing a sandwich-ELISA-based MSD immunoassay using the blood plasma collected one month post-injury. Both proteins are highly sensitive biomarkers that can indicate neurological injury in the clinical diagnosis of TBI ^59,60^. Two mice were removed due to failure to collect blood. We found that rmTBI was associated with a higher level of plasma Nf-L relative to the sham (*p*=0.0206; Fig. 6A), but that plasma GFAP was not different between rmTBI and sham at this post-injury time point (*p*=0.6730; Fig. 6B). These findings indicate that the CHIMERA rmTBI paradigm here induced enduring neurological injury in mice.

**Figure 6.**
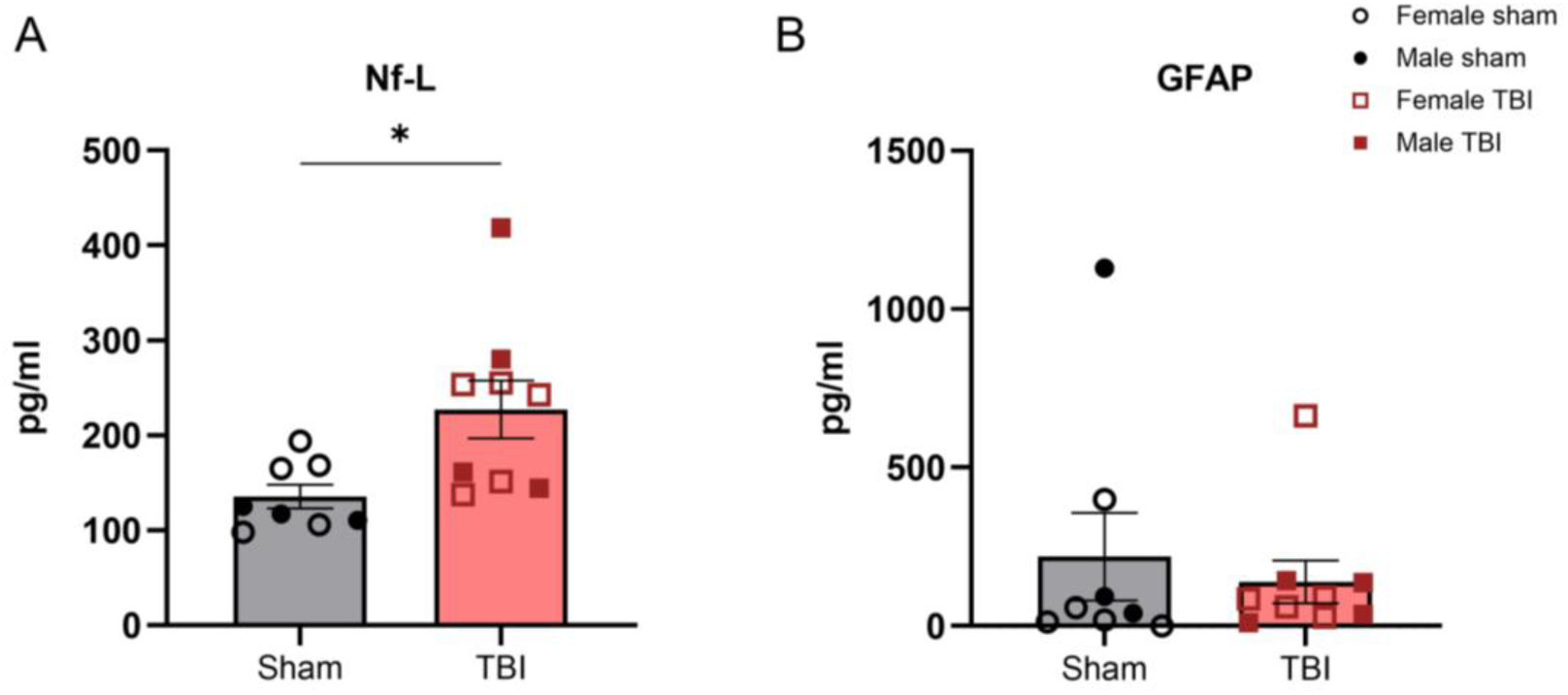
Plasma biomarkers after rmTBI. **(A)** Plasma Nf-L and **(B)** GFAP were compared between sham and TBI cohorts. Biomarker results were reported as mean ± SEM and analyzed by Mann Whitney-U test. Sham data are shown in black and rmTBI data are shown in red. Females are shown with open symbols and males are shown with solid symbols. Each data point represents a sample from a single mouse. Male TBI n=4, male sham n=3; female TBI n=5, female sham n=5. * p< 0.05.

### Amyloid histopathology was not exacerbated one month post-injury

To determine whether rmTBI exacerbates Aβ burden in the brain one month post-injury, we performed immunohistochemistry using the anti-Aβ antibody 6E10 to analyze Aβ histopathology. One female mouse was removed from histological analysis due to excessive tissue damage at cryosection. Overall, we found no differences in the level of amyloid plaques within the parietal isocortex and the hippocampus (Fig. 7A-C) after rmTBI, although amyloid pathology in females was significantly higher than in males in the isocortex (*p*=0.0177; Fig. 7B). We also analyzed regional amyloid pathology within several isolated cortical regions based on a published report ^50^, but again, we found no differences in the percentage area of amyloid plaques induced by rmTBI (Supplementary Fig. S5). There were also no group differences in the level of amyloid plaques in the anterior cortical regions (i.e., the site of injury; Supplementary Fig. S6).

**Figure 7.**
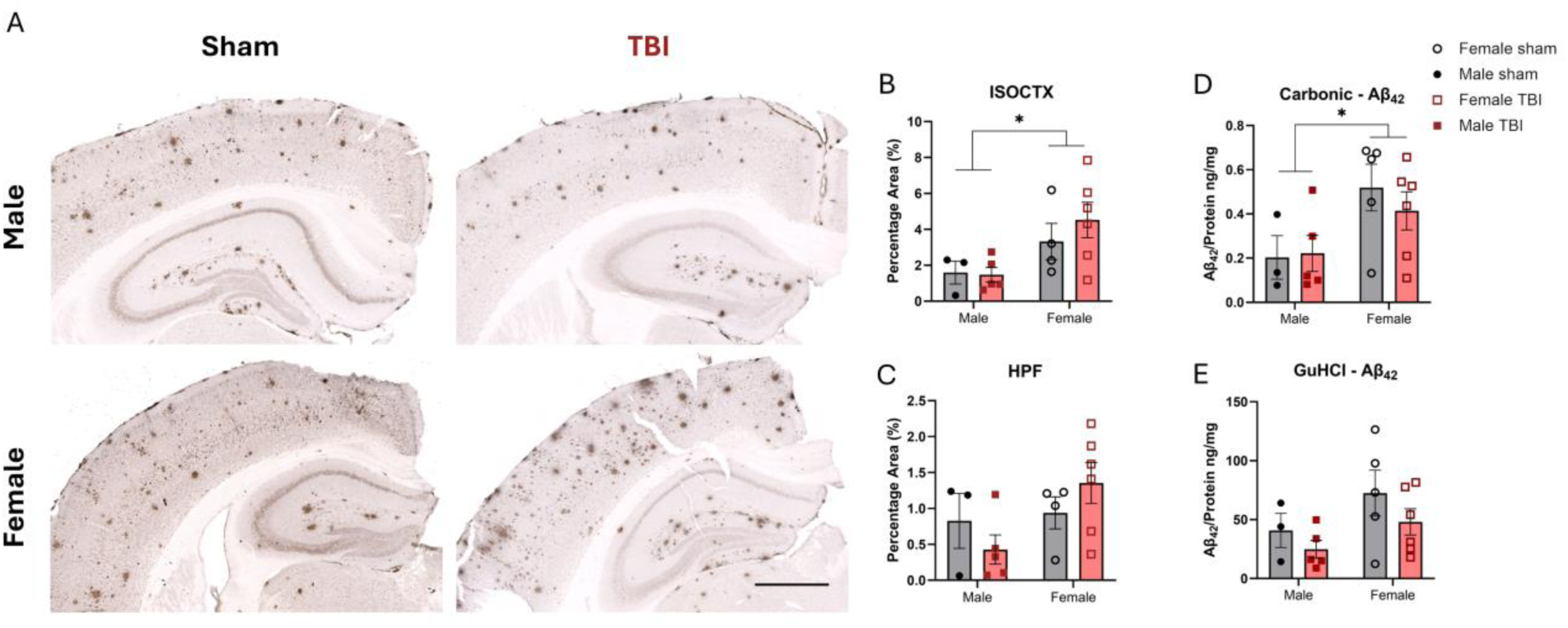
Amyloid pathology and Aβ42 after rmTBI. Amyloid plaques were immunostained by 6E10 one month post-injury. **(A)** Representative images of the isocortex and hippocampus of both male and female are shown. Scale bar = 1000 µm. Quantification of the percentage area of plaques in the **(B)** isocortex and **(C)** hippocampus of both sexes are shown on the right. Each data point represents the average of three sections per animal. Quantification of the **(D)** carbonic (soluble) and **(E)** GuHCl (insoluble) fractions of Aβ42 by ELISA using whole brain lysates. Each data point represents the average of three replicates per animal. Both histological and ELISA data are reported as mean ± SEM and analyzed by two-way ANOVA. Sham data are shown in black and rmTBI data are shown in red. Females are shown with open symbols and males are shown with solid symbols. For IHC, male TBI n=5, male sham n=3; female TBI n=6, female sham n=4. For ELISA, male TBI n=5, male sham n=3; female TBI n=6, female sham n=5. * *p*<0.05.

Additionally, we quantified the global level of soluble and insoluble monomeric Aβ_42_ in the carbonic and GuHCl fractions of the whole brain lysate by ELISA. Again, we did not observe significant differences in the level of Aβ_42_ (*p*>0.05) in both fractions, but a higher level of soluble Aβ_42_ (*p*=0.0202) and an uptrend in the level of insoluble Aβ_42_ (*p*=0.0744) were apparent in the female cohort regardless of injury (Fig 7 D-E), indicating a sex-dependent accumulation of Aβ_42_ monomer in the brain. Altogether, the neuropathological data suggested that rmTBI did not significantly alter amyloid pathology one month post-injury in the APP/PS1 mice.

## Discussion

Mild TBI is the most prevalent type of traumatic brain injury and is associated with increased risk of neurodegenerative disease, particularly when multiple injuries occur^61^. The mechanistic links between sleep disturbances, epileptiform activity, and neurodegenerative processes after TBI are not well understood. To explore the mechanisms underlying such changes, we characterized the effects of rmTBI on sleep, epileptiform activity, and Aβ pathology one month post-injury in a well-characterized mouse model of AD. We observed prolonged grimace signs after the second injury in female mice and evidence of neuronal damage in both sexes one month post-injury. RmTBI was also associated with a shift in the EEG power spectra during NREM sleep. However, we did not observe changes in vigilance state duration, epileptiform activity, or Aβ pathology at the time point examined. As expected, females exhibited higher amyloid burden and greater epileptiform activity than males.

Sleep disturbances have been linked to the earlier onset of mild cognitive impairments and increased Aβ burden ^62–65^. Sleep fragmentation has been associated with reduced slow-wave and REM sleep, slower recovery after TBI ^66^, and greater Aβ burden ^64^ in humans, and impaired glymphatic function in mice ^67^. In this study, we did not observe enduring changes in total sleep or sleep fragmentation measures, suggesting that sleep architecture could be relatively preserved in the APP/PS1 mice at one month post rmTBI. Prior animal studies have reported mixed effects of mTBI on sleep^17,68,69^, likely reflecting differences in injury model, injury severity, and post-injury time points. For example, Rowe et al. ^70^ reported that sleep changes following a midline fluid percussion injury normalize within one week, consistent with the absence of sleep disruptions at the one month time point in our cohort.

Spectral power analyses provide additional insight into brain activity across distinct frequency bands. During NREM sleep, lower frequencies typically dominate, reflecting increased neuronal synchrony, whereas higher frequencies are more prominent during wakefulness. Although sleep state duration did not differ between groups, rmTBI was associated with a shift toward higher frequencies during NREM, characterized by reduced low-frequency power and increased high-frequency power. The spectral shift was observed at the parietal electrode, consistent with the midline parietal impact site, and may indicate altered sleep quality in the rmTBI group.

Most quantitative EEG studies after TBI, in mice and humans, report increases in slow-wave power during wakefulness without major NREM alterations ^17,25,36^. However, studies on AD, excluding TBI, have observed different findings. In humans, decreased slow-wave activity has been associated with impaired Aβ clearance and cognitive deficits in older adults ^71^, as well as being observed in individuals diagnosed with mild to moderate AD ^71–73^. Similar spectral changes have been reported in AD mouse models ^39,46,62,74,75^. Thus, while NREM shifts are not typically observed after TBI alone, they are a recognized feature associated with AD pathology. In animal models, slow-wave activity has also been positively correlated with glymphatic influx ^34^, which is thought to support clearance of Aβ. Although we did not detect higher Aβ levels in the rmTBI group, these early spectral changes could potentially contribute to long-term impairments in brain clearance and increase Aβ burden.

Our findings suggest that rmTBI may further accentuate these slow-wave alterations in APP/PS1 mice, raising the possibility that reductions in slow-wave sleep after rmTBI reflect an interaction between the injury and underlying AD pathology. Whether such spectral shifts reflect early pathophysiological changes in sleep-related neural processes remains to be determined. Further work is needed to evaluate whether rmTBI impacts glymphatic function.

TBI has also been linked to post-traumatic epilepsy, through mechanisms including elevated iron levels, glutamate accumulation, impaired axonal transport, neurotransmitter imbalances, and epigenetic modifications that promote neuronal excitotoxicity ^76^. All of these changes lead to pathologic EEG activity such as high-frequency oscillations, abnormal sleep spindles, and epileptiform discharges ^76^. In this study, we assessed post-traumatic neuronal hyperactivity (spikes, SWDs, and seizures) but did not detect group differences at one month post-injury. This could be due to the chosen timing for EEG assessment, as prior work using severe TBI models has reported seizure onset beginning approximately 80 days post-injury ^77^. Thus, the one-month time point studied here may precede the period during which PTE events emerge. It is noteworthy that variability in epileptiform activity was greater in the sham group, potentially reflecting intrinsic characteristics of the APP/PS1 mouse, which exhibits elevated baseline epileptiform activity compared with wild-type mice ^40,78,79^. Researchers have proposed that overexpression of APP and, as a direct result, a premature increase in Aβ could be responsible for this increase in epileptiform activity ^20,78,80–82^. Female mice in our cohort displayed more SWDs than males, consistent with reports that sex can influence susceptibility to epileptiform activity. These observations underscore the importance of incorporating sex as a biological variable in future studies.

In our study, rmTBI was associated with elevated Nf-L levels in plasma, which is indicative of neurological injury one month post-injury. This aligns with clinical findings showing that increases in plasma Nf-L correlate with TBI severity^60^, and with preclinical findings demonstrating elevated Nf-L up to 5 weeks after closed-head mild-TBI ^10^. In contrast, we did not detect major differences in plasma GFAP between our sham and TBI cohorts at the investigated time point. Previous reports have shown increased plasma GFAP in wild-type mice 6 hours after a single high-energy CHIMERA impact ^49^. To our knowledge, GFAP responses to CHIMERA have not previously been evaluated in an amyloid-transgenic mouse model. Another study using a focal closed-head injury protocol reported elevated cortical GFAP in the APP/PS1 knock-in (KI) mouse model 2 months post-injury ^83^, suggesting that higher energy injuries can exacerbate astrogliosis in mice with existing amyloid pathology. In our study, the lack of GFAP response was perhaps due to the lower energy impacts used, as well as the small cohort size, particularly given that two sham mice exhibited unusually high baseline GFAP levels. This is also one of the first reports of plasma GFAP levels one month post-injury in a TBI animal model, as most preclinical studies have focused on acute windows ^84–87^. Clinical utility of plasma GFAP as a biomarker for TBI diagnosis appears limited to twelve months post-injury ^59,88–91^. Longitudinal profiling of plasma GFAP in both preclinical models and patients would provide a comprehensive outlook on how it may be utilized as a diagnostic tool for assessing chronic brain injury.

Proteopathic changes have been reported months after TBI in both clinical and preclinical studies. Positron emission tomography (PET) imaging with the ^11^C-PiB tracer suggested increased regional amyloid accumulation in patients one year after a moderate-to-severe TBI ^92^, and multiple studies in AD mouse modes have reported that TBI is associated with protein aggregation^93–95^. However, how repetitive mild TBI alter protein aggregation in mice remains unclear. We did not observe group differences in Aβ plaque burden at one month post-injury. Prior work suggests that the age of onset for amyloidosis in the transgenic APP mouse model may dictate the amyloidogenic outcome driven by TBI. For instance, reduced amyloid plaques below basal level have been reported in several transgenic APP mouse strains after receiving TBI at or beyond six months of age by using the controlled cortical impact^96^, the lateral fluid percussion injury^97^, and the CHIMERA^12^. This is perhaps due to changes in the clearance or enzymatic processing of Aβ after injury in older mice, where amyloid pathology has already been established. We also observed a high heterogeneity in the deposition of amyloid plaques at the investigated age in our cohort (Fig 7), indicating high variability in the basal level of amyloid pathology^43^. Based on these reports and our findings, we speculate that inducing rmTBI by CHIMERA in the APP/PS1 mice at younger ages prior to the onset of amyloidosis may manifest injury effects on amyloid pathology.

The lack of group differences across several outcome measures may reflect the relatively mild injury paradigm employed here, as well as the limited sample size. Any rmTBI-induced changes in sleep or neural activity may have been subtle, and the study may have been underpowered to detect these effects. Future studies could examine higher impact energies (0.7J), additional injuries (e.g. five or more), or modified inter-injury intervals to determine whether more robust or persistent effects emerge. We also acknowledge that male and female mice received a different number of injuries in this study, which may limit interpretation of sex effects. Although injury parameters were selected with the goal of achieving comparable levels of neurological injury, additional work is needed to establish sex-specific conditions that yield equivalent pathophysiological outcomes. Other limitations include methodological constraints, such as the need for single housing after headcap implantation, which may introduce variability due to stress. Although both groups were single-housed for the same duration, the effects of social isolation cannot be fully ruled out. Nevertheless, our study provides a feasible framework for the characterization of the electrophysiological, pathohistological, and plasma biomarkers following mTBI in a mouse model of AD. Continued refinement of injury parameters, timing, and multimodal outcome measures will be important for understanding how repetitive mild TBI interacts with AD-related processes across the lifespan.

## Acknowledgement

We thank Samantha Saw, Japneet Kaur, Bonnie K. Ng, Mayuko Arai and Robert Gibson for assisting in *in vivo* procedures and immunohistochemistry. We thank Dr. Sarah Faber for her insightful advice on power spectra analysis. We thank the SFU animal research facility for maintaining the animals used in this study.

## References

1. Lefevre-Dognin C, Cogné M, Perdrieau V, et al. Definition and epidemiology of mild traumatic brain injury. Neurochirurgie 2021;67(3):218–221; doi: 10.1016/j.neuchi.2020.02.002.

2. Van Den Heuvel C, Thornton E, Vink R. Traumatic Brain Injury and Alzheimer’s Disease: A Review. In: Progress in Brain Research. (Weber JT, Maas AIR. eds). Neurotrauma: New Insights into Pathology and Treatment Elsevier; 2007; pp. 303–316; doi: 10.1016/S0079-6123(06)61021-2.

3. Maas AIR, Menon DK, Manley GT, et al. Traumatic brain injury: progress and challenges in prevention, clinical care, and research. Lancet Neurol 2022;21(11):1004–1060; doi: 10.1016/S1474-4422(22)00309-X.

4. Thompson HJ, McCormick WC, Kagan SH. Traumatic Brain Injury in Older Adults: Epidemiology, Outcomes, and Future Implications. J Am Geriatr Soc 2006;54(10):1590–1595; doi: 10.1111/j.1532-5415.2006.00894.x.

5. Gardner RC, Burke JF, Nettiksimmons J, et al. Dementia Risk After Traumatic Brain Injury vs Nonbrain Trauma: The Role of Age and Severity. JAMA Neurol 2014;71(12):1490; doi: 10.1001/jamaneurol.2014.2668.

6. Huang YQ, Vyas MV, Bronskill SE, et al. Rate of incident dementia and care needs among older adults with new traumatic brain injury: a population-based cohort study. Can Med Assoc J 2025;197(33):E1067–E1077; doi: 10.1503/cmaj.250361.

7. Guskiewicz KM, McCrea M, Marshall SW, et al. Cumulative Effects Associated With Recurrent Concussion in Collegiate Football Players: The NCAA Concussion Study. JAMA 2003;290(19):2549; doi: 10.1001/jama.290.19.2549.

8. Montenigro PH, Alosco ML, Martin BM, et al. Cumulative Head Impact Exposure Predicts Later-Life Depression, Apathy, Executive Dysfunction, and Cognitive Impairment in Former High School and College Football Players. J Neurotrauma 2017;34(2):328–340; doi: 10.1089/neu.2016.4413.

9. Bugay V, Bozdemir E, Vigil FA, et al. A Mouse Model of Repetitive Blast Traumatic Brain Injury Reveals Post-Trauma Seizures and Increased Neuronal Excitability. J Neurotrauma 2020;37(2):248–261; doi: 10.1089/neu.2018.6333.

10. Moro F, Lisi I, Tolomeo D, et al. Acute Blood Levels of Neurofilament Light Indicate One-Year White Matter Pathology and Functional Impairment in Repetitive Mild Traumatic Brain Injured Mice. J Neurotrauma 2023;40(11–12):1144–1163; doi: 10.1089/neu.2022.0252.

11. Namjoshi DR, Cheng WH, McInnes KA, et al. Merging pathology with biomechanics using CHIMERA (Closed-Head Impact Model of Engineered Rotational Acceleration): a novel, surgery-free model of traumatic brain injury. Mol Neurodegener 2014;9(1):55; doi: 10.1186/1750-1326-9-55.

12. Cheng WH, Martens KM, Bashir A, et al. CHIMERA repetitive mild traumatic brain injury induces chronic behavioural and neuropathological phenotypes in wild-type and APP/PS1 mice. Alzheimers Res Ther 2019;11(1):6; doi: 10.1186/s13195-018-0461-0.

13. Cheng WH, Stukas S, Martens KM, et al. Age at injury and genotype modify acute inflammatory and neurofilament-light responses to mild CHIMERA traumatic brain injury in wild-type and APP/PS1 mice. Exp Neurol 2018;301:26–38; doi: 10.1016/j.expneurol.2017.12.007.

14. Conte V, Uryu K, Fujimoto S, et al. Vitamin E reduces amyloidosis and improves cognitive function in Tg2576 mice following repetitive concussive brain injury. J Neurochem 2004;90(3):758–764; doi: 10.1111/j.1471-4159.2004.02560.x.

15. Uryu K, Laurer H, McIntosh T, et al. Repetitive Mild Brain Trauma Accelerates Aβ Deposition, Lipid Peroxidation, and Cognitive Impairment in a Transgenic Mouse Model of Alzheimer Amyloidosis. J Neurosci 2002;22(2):446–454; doi: 10.1523/JNEUROSCI.22-02-00446.2002.

16. Petraglia AL, Plog BA, Dayawansa S, et al. The Spectrum of Neurobehavioral Sequelae after Repetitive Mild Traumatic Brain Injury: A Novel Mouse Model of Chronic Traumatic Encephalopathy. J Neurotrauma 2014;31(13):1211–1224; doi: 10.1089/neu.2013.3255.

17. Sandsmark DK, Elliott JE, Lim MM. Sleep-Wake Disturbances After Traumatic Brain Injury: Synthesis of Human and Animal Studies. Sleep 2017;40(5):zsx044; doi: 10.1093/sleep/zsx044.

18. Shandra O, Winemiller AR, Heithoff BP, et al. Repetitive Diffuse Mild Traumatic Brain Injury Causes an Atypical Astrocyte Response and Spontaneous Recurrent Seizures. J Neurosci 2019;39(10):1944–1963; doi: 10.1523/JNEUROSCI.1067-18.2018.

19. Kent BA, Feldman HH, Nygaard HB. Sleep and its regulation: An emerging pathogenic and treatment frontier in Alzheimer’s disease. Prog Neurobiol 2021;197:101902; doi: 10.1016/j.pneurobio.2020.101902.

20. Palop JJ, Chin J, Roberson ED, et al. Aberrant Excitatory Neuronal Activity and Compensatory Remodeling of Inhibitory Hippocampal Circuits in Mouse Models of Alzheimer’s Disease. Neuron 2007;55(5):697–711; doi: 10.1016/j.neuron.2007.07.025.

21. Vossel KA, Tartaglia MC, Nygaard HB, et al. Epileptic activity in Alzheimer’s disease: causes and clinical relevance. Lancet Neurol 2017;16(4):311–322; doi: 10.1016/S1474-4422(17)30044-3.

22. Aoun R, Rawal H, Attarian H, et al. Impact of traumatic brain injury on sleep: an overview. Nat Sci Sleep 2019;Volume 11:131–140; doi: 10.2147/NSS.S182158.

23. Morris AR, Gudenschwager Basso EK, Gutierrez-Monreal MA, et al. Lifelong Chronic Sleep Disruption in a Mouse Model of Traumatic Brain Injury. Neurotrauma Rep 2024;5(1):61–73; doi: 10.1089/neur.2023.0107.

24. Massimini M, Corbetta M, Sanchez-Vives MV, et al. Sleep-like cortical dynamics during wakefulness and their network effects following brain injury. Nat Commun 2024;15(1):7207; doi: 10.1038/s41467-024-51586-1.

25. Modarres MH, Kuzma NN, Kretzmer T, et al. EEG slow waves in traumatic brain injury: Convergent findings in mouse and man. Neurobiol Sleep Circadian Rhythms 2017;2:59–70; doi: 10.1016/j.nbscr.2016.06.001.

26. Nwakamma MC, Stillman AM, Gabard-Durnam LJ, et al. Slowing of Parameterized Resting-State Electroencephalography After Mild Traumatic Brain Injury. Neurotrauma Rep 2024;5(1):448–461; doi: 10.1089/neur.2024.0004.

27. Chikhi S, Matton N, Blanchet S. EEG power spectral measures of cognitive workload: A meta-analysis. Psychophysiology 2022;59(6); doi: 10.1111/psyp.14009.

28. Donoghue T, Haller M, Peterson EJ, et al. Parameterizing neural power spectra into periodic and aperiodic components. Nat Neurosci 2020;23(12):1655–1665; doi: 10.1038/s41593-020-00744-x.

29. Buzsáki G, Draguhn A. Neuronal Oscillations in Cortical Networks. Science 2004;304(5679):1926–1929; doi: 10.1126/science.1099745.

30. Flores-Sandoval AA, Davila-Pérez P, Buss SS, et al. Spectral power ratio as a measure of EEG changes in mild cognitive impairment due to Alzheimer’s disease: a case-control study. Neurobiol Aging 2023;130:50–60; doi: 10.1016/j.neurobiolaging.2023.05.010.

31. Iliff JJ, Chen MJ, Plog BA, et al. Impairment of Glymphatic Pathway Function Promotes Tau Pathology after Traumatic Brain Injury. J Neurosci 2014;34(49):16180–16193; doi: 10.1523/JNEUROSCI.3020-14.2014.

32. Jessen NA, Munk ASF, Lundgaard I, et al. The Glymphatic System: A Beginner’s Guide. Neurochem Res 2015;40(12):2583–2599; doi: 10.1007/s11064-015-1581-6.

33. Xie L, Kang H, Xu Q, et al. Sleep Drives Metabolite Clearance from the Adult Brain. Science 2013;342(6156):373–377; doi: 10.1126/science.1241224.

34. Hablitz LM, Vinitsky HS, Sun Q, et al. Increased glymphatic influx is correlated with high EEG delta power and low heart rate in mice under anesthesia. Sci Adv 2019;5(2):eaav5447; doi: 10.1126/sciadv.aav5447.

35. Dagum P, Giovangrandi L, Levendovszky SR, et al. A wireless device for continuous measurement of brain parenchymal resistance tracks glymphatic function in humans. Nat Biomed Eng 2025; doi: 10.1038/s41551-025-01394-9.

36. Konduru SS, Wallace EP, Pfammatter JA, et al. Sleep-wake characteristics in a mouse model of severe traumatic brain injury: Relation to posttraumatic epilepsy. Epilepsia Open 2021;6(1):181–194; doi: 10.1002/epi4.12462.

37. Saletti PG, Katsarou AM, Molero M, et al. Neurobiological Aspects of Post-Traumatic Epilepsy: Lessons from Animal Models. In: Post-Traumatic Epilepsy. (Mula M. ed) Cambridge University Press; 2021; pp. 1–28; doi: 10.1017/9781108644594.003.

38. Jankowsky JL, Slunt HH, Ratovitski T, et al. Co-expression of multiple transgenes in mouse CNS: a comparison of strategies. Biomol Eng 2001;17(6):157–165; doi: 10.1016/S1389-0344(01)00067-3.

39. Kent BA, Strittmatter SM, Nygaard HB. Sleep and EEG Power Spectral Analysis in Three Transgenic Mouse Models of Alzheimer’s Disease: APP/PS1, 3xTgAD, and Tg2576. J Alzheimers Dis 2018;64(4):1325–1336; doi: 10.3233/JAD-180260.

40. Gureviciene I, Ishchenko I, Ziyatdinova S, et al. Characterization of Epileptic Spiking Associated With Brain Amyloidosis in APP/PS1 Mice. Front Neurol 2019;10:1151; doi: 10.3389/fneur.2019.01151.

41. Nygaard HB, Kaufman AC, Sekine-Konno T, et al. Brivaracetam, but not ethosuximide, reverses memory impairments in an Alzheimer’s disease mouse model. Alzheimers Res Ther 2015;7(1):25; doi: 10.1186/s13195-015-0110-9.

42. Namjoshi DR, Cheng WH, Bashir A, et al. Defining the biomechanical and biological threshold of murine mild traumatic brain injury using CHIMERA (Closed Head Impact Model of Engineered Rotational Acceleration). Exp Neurol 2017;292:80–91; doi: 10.1016/j.expneurol.2017.03.003.

43. Yan P, Bero AW, Cirrito JR, et al. Characterizing the Appearance and Growth of Amyloid Plaques in APP/PS1 Mice. J Neurosci 2009;29(34):10706–10714; doi: 10.1523/JNEUROSCI.2637-09.2009.

44. Garcia-Alloza M, Robbins EM, Zhang-Nunes SX, et al. Characterization of amyloid deposition in the APPswe/PS1dE9 mouse model of Alzheimer disease. Neurobiol Dis 2006;24(3):516–524; doi: 10.1016/j.nbd.2006.08.017.

45. Langford DJ, Bailey AL, Chanda ML, et al. Coding of facial expressions of pain in the laboratory mouse. Nat Methods 2010;7(6):447–449; doi: 10.1038/nmeth.1455.

46. Kent BA, Michalik M, Marchant EG, et al. Delayed daily activity and reduced NREM slow-wave power in the APPswe/PS1dE9 mouse model of Alzheimer’s disease. Neurobiol Aging 2019;78:74–86; doi: 10.1016/j.neurobiolaging.2019.01.010.

47. Kadam SD, D’Ambrosio R, Duveau V, et al. Methodological standards and interpretation of video-electroencephalography in adult control rodents. A TASK1-WG1 report of theAES/ILAETranslational Task Force of the ILAE. Epilepsia 2017;58(S4):10–27; doi: 10.1111/epi.13903.

48. Jin N, Babiloni C, Drinkenburg WH, et al. Recommendations for Preclinical Testing of Treatments Against Alzheimer’s Disease-Related Epileptiform Spikes in Transgenic Rodent Models. J Alzheimers Dis JAD 2022;88(3):849–865; doi: 10.3233/JAD-210209.

49. Button EB, Cheng WH, Barron C, et al. Development of a novel, sensitive translational immunoassay to detect plasma glial fibrillary acidic protein (GFAP) after murine traumatic brain injury. Alzheimers Res Ther 2021;13(1):58; doi: 10.1186/s13195-021-00793-9.

50. Vasciaveo V, Iadarola A, Casile A, et al. Sleep fragmentation affects glymphatic system through the different expression of AQP4 in wild type and 5xFAD mouse models. Acta Neuropathol Commun 2023;11(1):16; doi: 10.1186/s40478-022-01498-2.

51. Palmqvist S, Schöll M, Strandberg O, et al. Earliest accumulation of β-amyloid occurs within the default-mode network and concurrently affects brain connectivity. Nat Commun 2017;8(1):1214; doi: 10.1038/s41467-017-01150-x.

52. Murphy C. Loss of Olfactory Function in Dementing Disease. Physiol Behav 1999;66(2):177–182; doi: 10.1016/S0031-9384(98)00262-5.

53. Mishra P, Pandey C, Singh U, et al. Descriptive statistics and normality tests for statistical data. Ann Card Anaesth 2019;22(1):67; doi: 10.4103/aca.ACA_157_18.

54. McIntosh AR, Lobaugh NJ. Partial least squares analysis of neuroimaging data: applications and advances. NeuroImage 2004;23:S250–S263; doi: 10.1016/j.neuroimage.2004.07.020.

55. Cragg L, Kovacevic N, McIntosh AR, et al. Maturation of EEG power spectra in early adolescence: a longitudinal study: Maturation of EEG. Dev Sci 2011;14(5):935–943; doi: 10.1111/j.1467-7687.2010.01031.x.

56. Wang J, Tanila H, Puoliväli J, et al. Gender differences in the amount and deposition of amyloidβ in APPswe and PS1 double transgenic mice. Neurobiol Dis 2003;14(3):318–327; doi: 10.1016/j.nbd.2003.08.009.

57. Berman R, Spencer H, Boese M, et al. Loss of Consciousness and Righting Reflex Following Traumatic Brain Injury: Predictors of Post-Injury Symptom Development (A Narrative Review). Brain Sci 2023;13(5):750; doi: 10.3390/brainsci13050750.

58. Tok S, Crauwels D, Drinkenburg W. A Characterization of Epileptiform Activity Associated with TAU Seeding and Amyloid Pathology in the APPKM670/671NL.PS1/L166P and APP-KI Animal Models. J Neurol Exp Neurosci 2022;08(01); doi: 10.17756/jnen.2022-096.

59. Graham NSN, Zimmerman KA, Moro F, et al. Axonal marker neurofilament light predicts long-term outcomes and progressive neurodegeneration after traumatic brain injury. Sci Transl Med 2021;13(613):eabg9922; doi: 10.1126/scitranslmed.abg9922.

60. Shahim P, Politis A, Van Der Merwe A, et al. Neurofilament light as a biomarker in traumatic brain injury. Neurology 2020;95(6); doi: 10.1212/WNL.0000000000009983.

61. Wilson L, Stewart W, Dams-O’Connor K, et al. The chronic and evolving neurological consequences of traumatic brain injury. Lancet Neurol 2017;16(10):813–825; doi: 10.1016/S1474-4422(17)30279-X.

62. Holth JK, Mahan TE, Robinson GO, et al. Altered sleep and EEG power in the P301S Tau transgenic mouse model. Ann Clin Transl Neurol 2017;4(3):180–190; doi: 10.1002/acn3.390.

63. Mander BA, Winer JR, Jagust WJ, et al. Sleep: A Novel Mechanistic Pathway, Biomarker, and Treatment Target in the Pathology of Alzheimer’s Disease? Trends Neurosci 2016;39(8):552–566; doi: 10.1016/j.tins.2016.05.002.

64. Nguyen Ho PT, Hoepel SJW, Rodriguez-Ayllon M, et al. Sleep, 24-Hour Activity Rhythms, and Subsequent Amyloid-β Pathology. JAMA Neurol 2024;81(8):824–834; doi: 10.1001/jamaneurol.2024.1755.

65. Roh JH, Huang Y, Bero AW, et al. Disruption of the Sleep-Wake Cycle and Diurnal Fluctuation of β-Amyloid in Mice with Alzheimer’s Disease Pathology. Sci Transl Med 2012;4(150):150ra122–150ra122; doi: 10.1126/scitranslmed.3004291.

66. Fleming MK, Smejka T, Henderson Slater D, et al. Sleep Disruption After Brain Injury Is Associated With Worse Motor Outcomes and Slower Functional Recovery. Neurorehabil Neural Repair 2020;34(7):661–671; doi: 10.1177/1545968320929669.

67. Deng S, Hu Y, Chen S, et al. Chronic sleep fragmentation impairs brain interstitial clearance in young wildtype mice. J Cereb Blood Flow Metab 2024;44(9):1515–1531; doi: 10.1177/0271678X241230188.

68. Korthas HT, Main BS, Harvey AC, et al. The Effect of Traumatic Brain Injury on Sleep Architecture and Circadian Rhythms in Mice—A Comparison of High-Frequency Head Impact and Controlled Cortical Injury. Biology 2022;11(7):1031; doi: 10.3390/biology11071031.

69. Portillo E, Zi X, Kim Y, et al. Persistent hypersomnia following repetitive mild experimental traumatic brain injury: Roles of chronic stress and sex differences. J Neurosci Res 2023;101(6):843–865; doi: 10.1002/jnr.25165.

70. Rowe RK, Harrison JL, O’Hara BF, et al. Diffuse brain injury does not affect chronic sleep patterns in the mouse. Brain Inj 2014;28(4):504–510; doi: 10.3109/02699052.2014.888768.

71. Liguori C, Placidi F, Izzi F, et al. Sleep dysregulation, memory impairment, and CSF biomarkers during different levels of neurocognitive functioning in Alzheimer’s disease course. Alzheimers Res Ther 2020;12(1):5; doi: 10.1186/s13195-019-0571-3.

72. Kent BA, Casciola AA, Carlucci SK, et al. Home EEG sleep assessment shows reduced slow-wave sleep in mild–moderate Alzheimer’s disease. Alzheimers Dement Transl Res Clin Interv 2022;8(1):e12347; doi: 10.1002/trc2.12347.

73. Liu S, Pan J, Tang K, et al. Sleep spindles, K-complexes, limb movements and sleep stage proportions may be biomarkers for amnestic mild cognitive impairment and Alzheimer’s disease. Sleep Breath 2020;24(2):637–651; doi: 10.1007/s11325-019-01970-9.

74. Chen C-W, Kwok Y-T, Cheng Y-T, et al. Reduced slow-wave activity and autonomic dysfunction during sleep precede cognitive deficits in Alzheimer’s disease transgenic mice. Sci Rep 2023;13(1); doi: 10.1038/s41598-023-38214-6.

75. Zhang F, Zhong R, Li S, et al. Alteration in sleep architecture and electroencephalogram as an early sign of Alzheimer’s disease preceding the disease pathology and cognitive decline. Alzheimers Dement 2019;15(4):590–597; doi: 10.1016/j.jalz.2018.12.004.

76. Golub VM, Reddy DS. Post-Traumatic Epilepsy and Comorbidities: Advanced Models, Molecular Mechanisms, Biomarkers, and Novel Therapeutic Interventions. France C. ed. Pharmacol Rev 2022;74(2):387–438; doi: 10.1124/pharmrev.121.000375.

77. Guo D, Zeng L, Brody DL, et al. Rapamycin Attenuates the Development of Posttraumatic Epilepsy in a Mouse Model of Traumatic Brain Injury. Borlongan CV. ed. PLoS ONE 2013;8(5):e64078; doi: 10.1371/journal.pone.0064078.

78. Minkeviciene R, Rheims S, Dobszay MB, et al. Amyloid β-Induced Neuronal Hyperexcitability Triggers Progressive Epilepsy. J Neurosci 2009;29(11):3453–3462; doi: 10.1523/JNEUROSCI.5215-08.2009.

79. Reyes-Marin KE, Nuñez A. Seizure susceptibility in the APP/PS1 mouse model of Alzheimer’s disease and relationship with amyloid β plaques. Brain Res 2017;1677:93–100; doi: 10.1016/j.brainres.2017.09.026.

80. Hijazi S, Smit AB, Van Kesteren RE. Fast-spiking parvalbumin-positive interneurons in brain physiology and Alzheimer’s disease. Mol Psychiatry 2023;28(12):4954–4967; doi: 10.1038/s41380-023-02168-y.

81. Tok S, Ahnaou A, Drinkenburg W. Functional Neurophysiological Biomarkers of Early-Stage Alzheimer’s Disease: A Perspective of Network Hyperexcitability in Disease Progression. J Alzheimers Dis 2022;88(3):809–836; doi: 10.3233/jad-210397.

82. Verret L, Mann EO, Hang GB, et al. Inhibitory Interneuron Deficit Links Altered Network Activity and Cognitive Dysfunction in Alzheimer Model. Cell 2012;149(3):708–721; doi: 10.1016/j.cell.2012.02.046.

83. Webster SJ, Van Eldik LJ, Watterson DM, et al. Closed Head Injury in an Age-Related Alzheimer Mouse Model Leads to an Altered Neuroinflammatory Response and Persistent Cognitive Impairment. J Neurosci 2015;35(16):6554–6569; doi: 10.1523/JNEUROSCI.0291-15.2015.

84. Cikriklar HI, Onur U, Ekici MA, et al. Effectiveness of gfap in determining neuron damage in rats with induced head trauma. Turk Neurosurg 2015; doi: 10.5137/1019-5149.JTN.13946-15.2.

85. Kmeťová K, Drobná D, Lipták R, et al. Early dynamics of glial fibrillary acidic protein and extracellular DNA in plasma of mice after closed head traumatic brain injury. Neurochirurgie 2022;68(6):e68–e74; doi: 10.1016/j.neuchi.2022.06.003.

86. Lafrenaye AD, Mondello S, Wang KK, et al. Circulating GFAP and Iba-1 levels are associated with pathophysiological sequelae in the thalamus in a pig model of mild TBI. Sci Rep 2020;10(1):13369; doi: 10.1038/s41598-020-70266-w.

87. Wei C, Luo Y, Peng L, et al. Expression of Notch and Wnt/β-catenin signaling pathway in acute phase severe brain injury rats and the effect of exogenous thyroxine on those pathways. Eur J Trauma Emerg Surg 2021;47(6):2001–2015; doi: 10.1007/s00068-020-01359-4.

88. Bogoslovsky T, Wilson D, Chen Y, et al. Increases of Plasma Levels of Glial Fibrillary Acidic Protein, Tau, and Amyloid β up to 90 Days after Traumatic Brain Injury. J Neurotrauma 2017;34(1):66–73; doi: 10.1089/neu.2015.4333.

89. Gardner RC, Puccio AM, Korley FK, et al. Effects of age and time since injury on traumatic brain injury blood biomarkers: a TRACK-TBI study. Brain Commun 2022;5(1):fcac316; doi: 10.1093/braincomms/fcac316.

90. Metting Z, Wilczak N, Rodiger LA, et al. GFAP and S100B in the acute phase of mild traumatic brain injury. Neurology 2012;78(18):1428–1433; doi: 10.1212/WNL.0b013e318253d5c7.

91. Papa L, Silvestri S, Brophy GM, et al. GFAP Out-Performs S100β in Detecting Traumatic Intracranial Lesions on Computed Tomography in Trauma Patients with Mild Traumatic Brain Injury and Those with Extracranial Lesions. J Neurotrauma 2014;31(22):1815–1822; doi: 10.1089/neu.2013.3245.

92. Scott G, Ramlackhansingh AF, Edison P, et al. Amyloid pathology and axonal injury after brain trauma. Neurology 2016;86(9):821–828; doi: 10.1212/WNL.0000000000002413.

93. Alyenbaawi H, Kanyo R, Locskai LF, et al. Seizures are a druggable mechanistic link between TBI and subsequent tauopathy. eLife 2021;10:e58744; doi: 10.7554/eLife.58744.

94. Edwards G, Zhao J, Dash PK, et al. Traumatic Brain Injury Induces Tau Aggregation and Spreading. J Neurotrauma 2020;37(1):80–92; doi: 10.1089/neu.2018.6348.

95. Washington PM, Morffy N, Parsadanian M, et al. Experimental Traumatic Brain Injury Induces Rapid Aggregation and Oligomerization of Amyloid-Beta in an Alzheimer’s Disease Mouse Model. J Neurotrauma 2014;31(1):125–134; doi: 10.1089/neu.2013.3017.

96. Nakagawa Y, Reed L, Nakamura M, et al. Brain Trauma in Aged Transgenic Mice Induces Regression of Established Aβ Deposits. Exp Neurol 2000;163(1):244–252; doi: 10.1006/exnr.2000.7375.

97. Collins JM, King AE, Woodhouse A, et al. Age Moderates the Effects of Traumatic Brain Injury on Beta-Amyloid Plaque Load in APP/PS1 Mice. J Neurotrauma 2019;36(11):1876–1889; doi: 10.1089/neu.2018.5982.

