## Supplementary figures for "Repetitive mild traumatic brain injury causes neuronal damage in the APP/PS1 mouse model of Alzheimer’s disease without an enduring impact on amyloid pathology, sleep, or epileptiform activity"

**Figure S1**

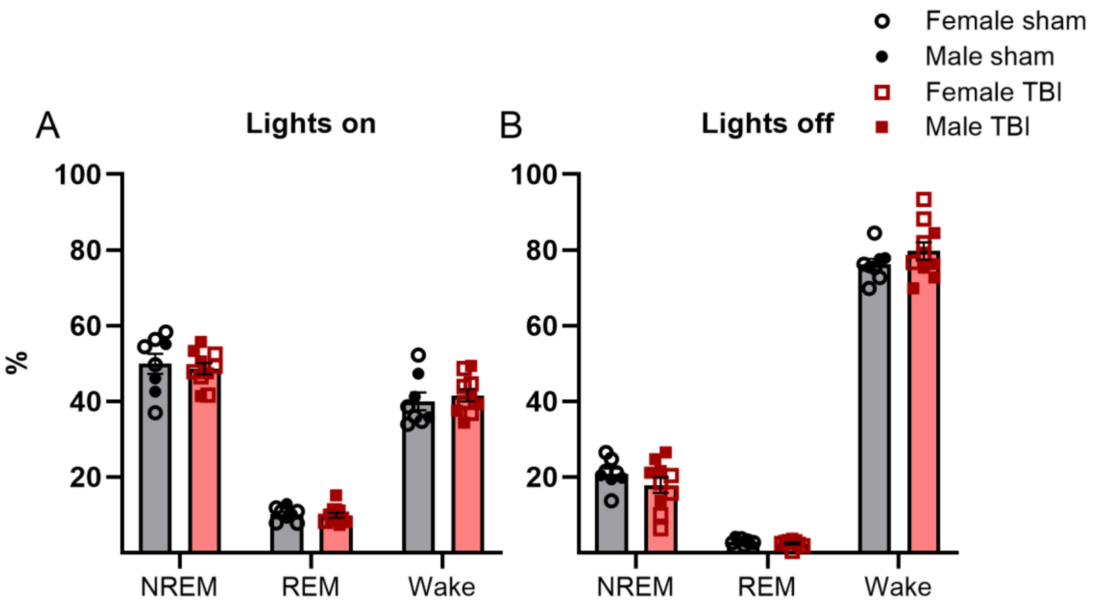


**Figure S1. Percentage of time spent in each vigilance state during lights on and off periods**. There were no statistically significant differences between the TBI and sham groups in the amount of time spent at each vigilance state during lights-on **(A)** or lights-off **(B)** periods. All data are expressed in mean ±SEM and analyzed by Mann Whitney-U test. Sham data are shown in black and rmTBI data are shown in red. Females are shown with open symbols and males are shown with solid symbols. Male TBI n=5, male sham n=3; female TBI n=5, female sham n=5.

**Figure S2.**

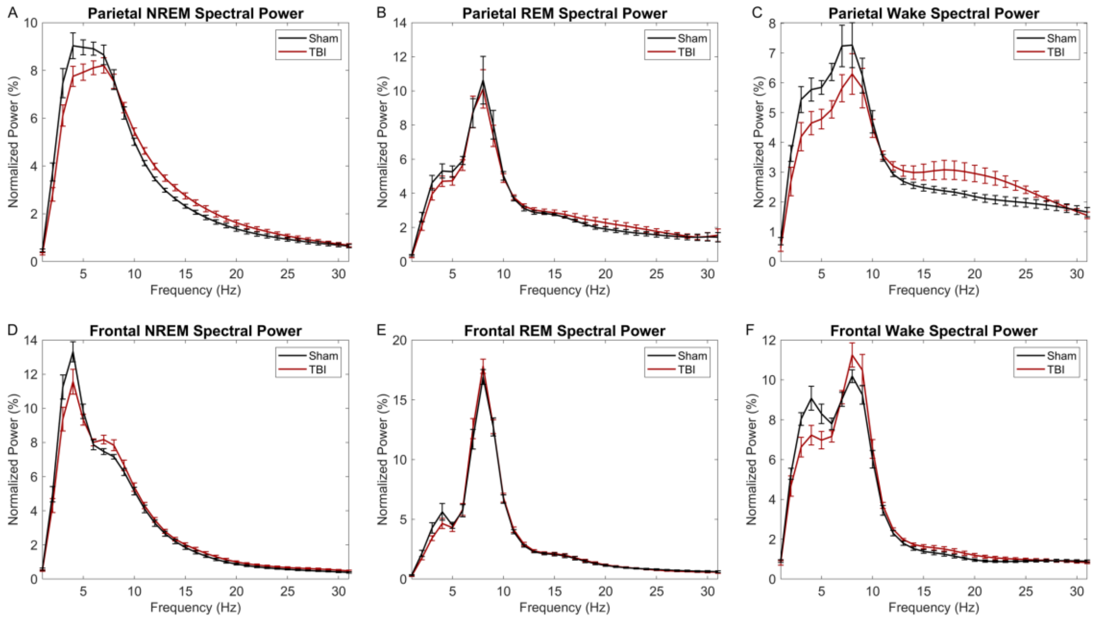


**Figure S2. Frontal and parietal spectral power graphs across vigilance states**. Normalized spectral power graphs for parietal NREM **(A)**, REM **(B)**, and wake **(C)** and frontal NREM **(D)**, REM **(E)**, and wake **(F)** across the 24h recording for frequencies 0.5-30Hz. The spectral power for parietal NREM, previously presented in figure 4, is also presented here to allow for direct comparison. Partial least squares analysis (PLS) found no differences between the groups at either of these states or locations, with the exception of the parietal NREM spectral power results presented before. Sex differences were not evaluated due to sample size limitations. Error bars represent the ±SEM and data was analyzed using the PLS method. Sham data is shown in black and the rmTBI data is shown in red. Male TBI n=5, male sham n=3; female TBI n=5, female sham n=5.

**Figure S3.**

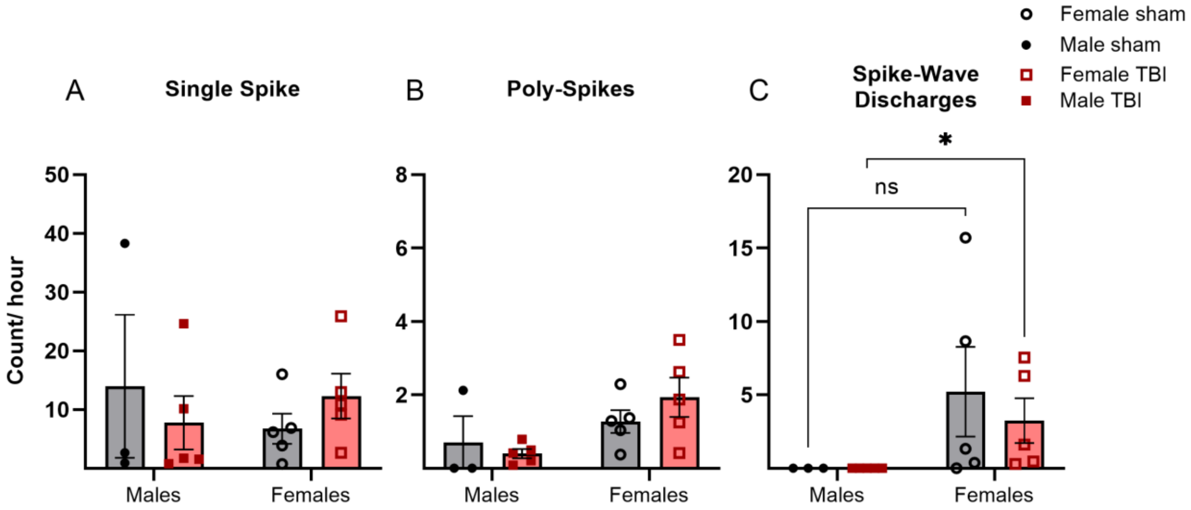


**Figure S3. Epileptiform activity one month after rmTBI.** There were no sex or TBI effects on **(A)** single spike or **(B)** poly spikes counts per hour after rmTBI one month post-injury. **(C)** Spike-wave discharges (SWDs) were only observed in female mice. Female TBI mice had significantly more SWDs counts per hour compared to male TBI mice, there were no sex differences within the sham group. Data are reported as mean ± SEM and analyzed by Kruskal-Wallis with Dunn's correction. Sham data are shown in black and rmTBI data are shown in red. Females are shown with open symbols and males are shown with solid symbols. rmTBI n=10, sham n=8; males n=8, females n=10. Male TBI n=5, male sham n=3; female TBI n=5, female sham n=5. * $p<0.05$.

**Figure S4.**

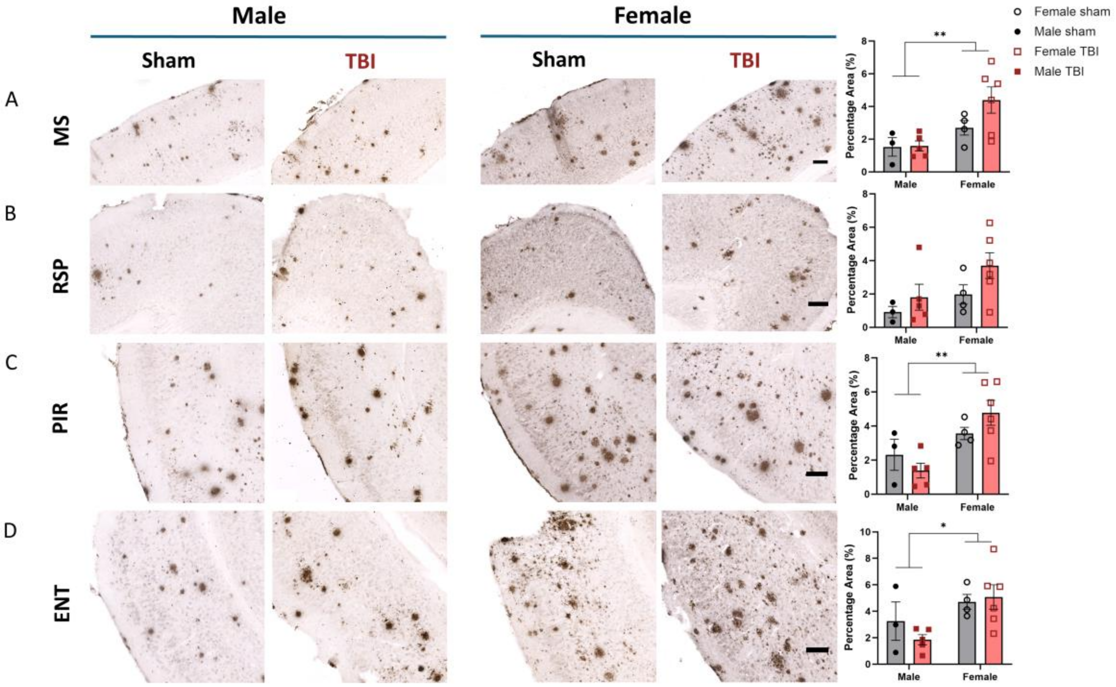

**Figure S4. Regional amyloid histopathology in the parietal cortex after rmTBI.** Representative images of 6E10 immunostained brain sections in the **(A)** motor sensory cortex (MS), **(B)** retrosplenial cortex (RSP), **(C)** piriform area (PIR) and **(D)** entorhinal cortex (ENT). Quantification plots of the percentage area of 6E10 positive plaque within the defined region of interest are shown on the right. Each data point represents the average of three sections per animal. Data are reported as mean ± SEM and analyzed by two-way ANOVA. Scale bar = 200 µm. Sham data are shown in black and rmTBI data are shown in red. Females are shown with open symbols and males are shown with solid symbols. Male TBI n=5, male sham n=3; female TBI n=6, female sham n=4. * p<0.05. ** p<0.01.

**Figure S5.**

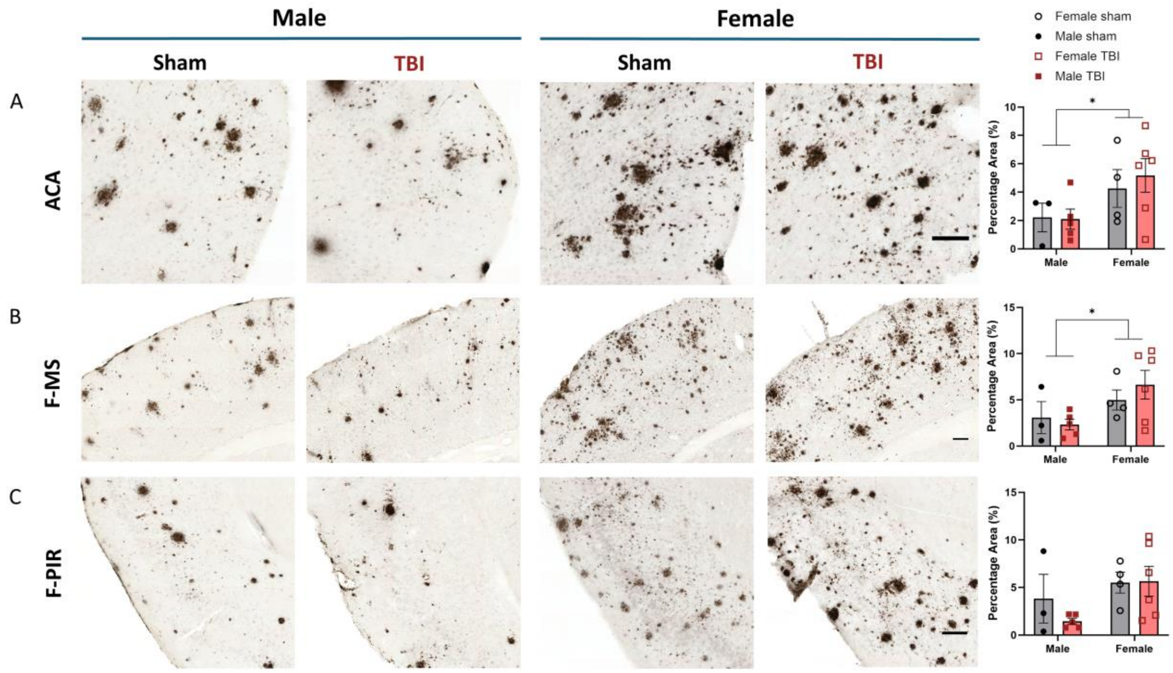


**Figure S5. Amyloid histopathology in the anterior cortical regions after rmTBI.** Representative images of 6E10 immunostained brain sections in the (**A**) anterior cingulate area (ACA), (**B**) frontal motor sensory cortex (F-MS), and (**C**) frontal piriform area (F-PIR). Quantification plots of the percentage area of 6E10 positive plaque within the defined region of interest are shown on the right. Each data point represents the average of three sections per animal. Data are reported as mean ± SEM and analyzed by two-way ANOVA. Scale bar = 100 µm. Sham data are shown in black and rmTBI data are shown in red. Females are shown with open symbols and males are shown with solid symbols. Male TBI n=5, male sham n=3; female TBI n=6, female sham n=4. * p<0.05.
